# Beyond Histones: Unveiling the Functional Roles of Protein Acetylation in Prokaryotes and Eukaryotes

**DOI:** 10.1101/2024.06.26.600871

**Authors:** Bruno Sousa Bonifácio, Ariely Barbosa Leite, Ana Caroline de Castro Nascimento Sousa, Suellen Rodrigues Maran, Antoniel Augusto Severo Gomes, Elton J. R. Vasconcelos, Nilmar Silvio Moretti

**Author notes:** Corresponding: Nilmar Silvio Moretti.

## Abstract

Lysine acetylation plays a crucial role in cellular processes and is found across various evolutionary organisms. Recent advancements in proteomic techniques revealed the presence of acetylation in thousands of non-histone proteins. Here, we conducted extensive meta-analysis of 48 acetylomes spanning diverse organisms, including archaea, bacteria, fungi, protozoa, worms, plants, insects, crustacea, fish, and mammals. Our analyzes revealed a predominance of a single acetylation site in a protein detected in all studied organisms, and proteins heavily acetylated, with >5-10 acetylated-sites, were represented by Hsp70, histone or transcription GTP-biding domain. Moreover, using gene enrichment approaches we found that ATP metabolic processes, glycolysis, aminoacyl-tRNA synthetase pathways and oxidative stress response are among the most acetylated cellular processes. Finally, to better explore the regulatory function of acetylation in glycolysis and oxidative stress we used aldolase and superoxide dismutase A (SODA) enzymes as model. For aldolase, we found that K147 acetylation, responsible to regulate human enzyme, conserved in all phylogenic clade, suggesting that this acetylation might play the same role in other species; while for SODA, we identified many lysine residues in different species present in the tunnel region, which was demonstrated for human and *Trypanosoma cruzi,* as negative regulator, also suggesting a conserved regulatory mechanism. In conclusion, this study provides insights into the conservation and functional significance of lysine acetylation in different organisms emphasizing its roles in cellular processes, metabolic pathways, and molecular regulation, shedding light in the extensive function of non-histone lysine acetylation.

## INTRODUCTION

The ability of cells to detect and respond appropriately to their surrounding environment is essential for the functioning of every living organism. Cells are constantly exposed to a range of different stimuli; they can adapt to each new situation due to the existence of intracellular signaling pathways. Cellular processes that allow cells to adapt to these situations can be regulated through reversible post-translational modifications of lipids, nucleic acids, and proteins. Hundreds of protein post-translational modifications have been described, such as methylation, phosphorylation, crotonylation, and acetylation [1–3].

Protein acetylation was first described in 1963 in the N-terminal region of histone [4]. For decades, research focused on lysine acetylation in histone proteins, particularly how this modification impacts chromatin structure and gene expression. However, the development of more accurate and sensitive proteomic techniques has enabled the identification of thousands of acetylated non-histone proteins across various prokaryotic and eukaryotic species [5–7].

The set of lysine-acetylated proteins of an organism is known as acetylome. The first acetylome, published in 2009 by Choudhary et al., [8] described the Kac proteins of HeLa cells. Since then, researchers have characterized the acetylomes of various organisms including *Saccharomyces cerevisiae, Escherichia coli, Drosophila melanogaster, Toxoplasma gondii, Plasmodium falciparum, Schistosoma japonicum*, and *Homo sapiens* [6, 9–14]. These global proteomic analyses revealed a vast array of proteins acetylated across diverse cellular compartments, suggesting their involvement in a multitude of cellular functions, including metabolism, translation, response to oxidative stress, and gene expression regulation [6, 7].

Lysine acetylation, which can occur enzymatically or spontaneously, neutralizes the positive charge of the lysine, potentially affecting enzymatic activity, protein-protein interactions, and subcellular localization [5, 7]. This highlights the importance of lysine acetylation in regulating cellular processes, emerging as a key player alongside phosphorylation in this vital role.

While a significant number of acetylomes have been published, a gap remains in our understanding of the specific regulatory roles of lysine acetylation across diverse organisms. To address this, we performed extensive analyses of 50 acetylomes spanning bacteria, fungi, protozoa, worms, plants insects, fish, and mammals. Our findings suggest a potential for conserved roles of lysine acetylation in specific protein groups, particularly heat-shock proteins, glycolysis/TCA cycle enzymes, and antioxidant enzymes. This work sheds light on the potential mechanisms of protein acetylation and its impact on core cellular processes across different kingdoms.

## RESULTS

### Prokaryotes and eukaryotes exhibit extensive lysine acetylation in non-histone proteins

To gauge the extent of our knowledge of non-histone protein acetylation, we conducted a comprehensive literature search for published studies describing prokaryotic and eukaryotic acetylomes from 2009 (following the first published acetylome) to 2022. This search identified approximately 128 articles (Figure 1A) spanning diverse organisms, including archaea, bacteria, fungi, protozoa, worms, plants, insects, crustacea, fish, and mammals (Figure 1B). Notably, bacteria dominated the research landscape, likely due to a single study characterizing the acetylome of 48 bacterial species [15].

**Figure 1.**
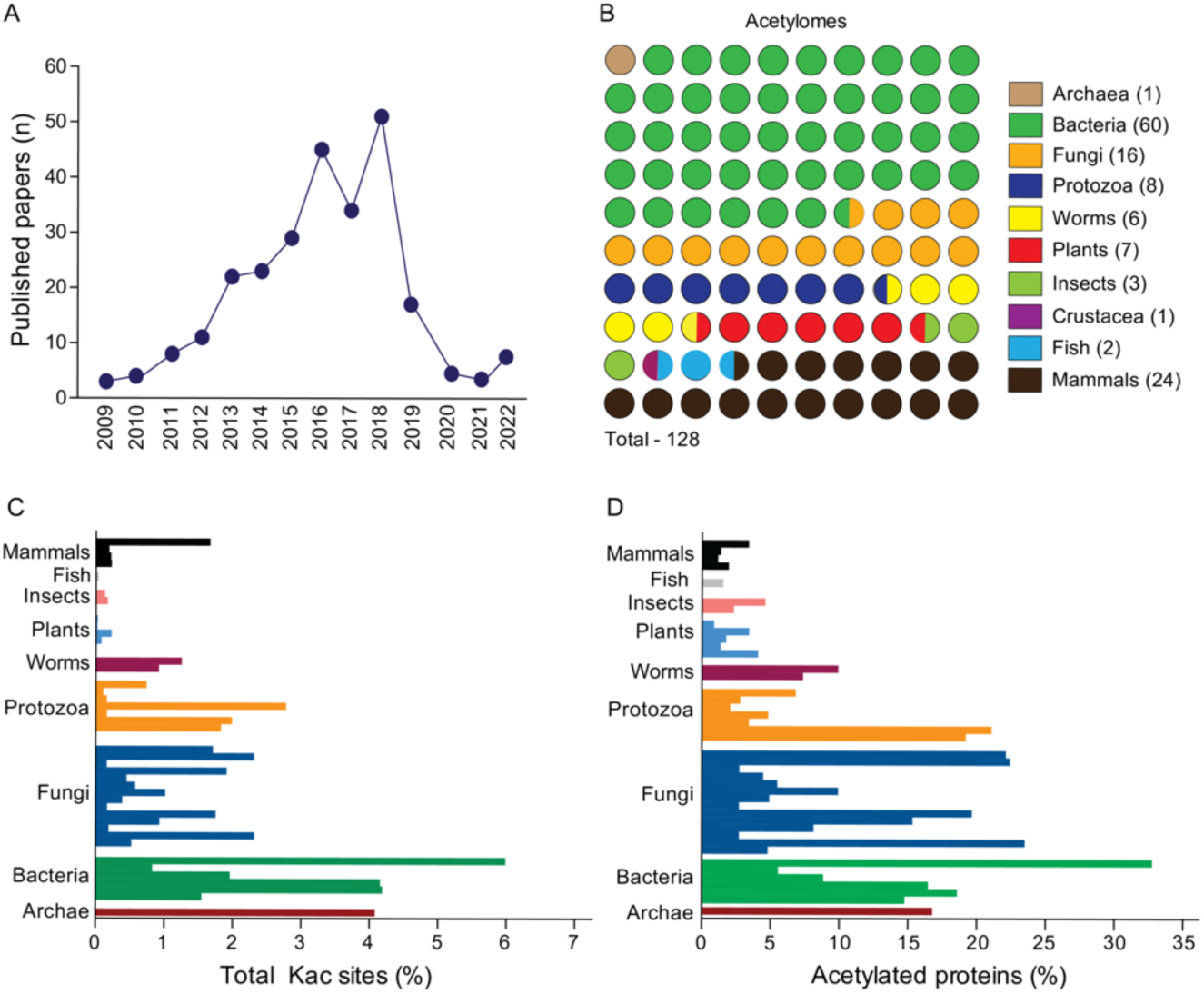
Lysine acetylation is widely detected in different groups of organisms. **A.** Number of articles published describing acetylomes from 2009-2022. **B.** Distribution of published acetylomes based on groups of organisms (archaea, bacteria, fungi, protozoa, worms, plants, insects, crustacea, fish and mammals). **C.** Percentage of total lysine-acetylated sites found based on total lysine residues of each proteome and total lysine-acetylated detected in each acetylome. **D.** Percentage of total acetylated proteins found based on proteome size and lysine-acetylated protein identified per acetylome. Each bar represents the acetylome of a specific species and for more details about each specie see Figure Sup 1.

To ensure data consistency for further analysis, we selected only studies employing lysine-acetylated peptide enrichment prior to mass spectrometry. Additionally, we excluded crustaceans due to the lack of a published genome for *Alvinocaris longirostris* (deep sea shrimp), the only species with a described acetylome during our search period. This resulted in a dataset encompassing 48 acetylomes from 37 species distributed across various taxa: Archaea (1), Bacteria (6), Fungi (15), Protozoa (8), Worms (6), Plants (5), Insects (2), Fish (1), and Mammals (4) (Table S1, Figure S1).

Based on the selected acetylomes and the number of lysine residues and proteins within each species’ proteome, we calculated the percentage of both total lysine-acetylated residues (Kac sites) and total acetylated proteins (Kac proteins) for each species. Overall, the percentage of Kac sites ranged from 0.15% to 5.99%, with an average of around 2.5% (Figures 1C and S2). *Vibrio cholerae* displayed the highest percentage of Kac sites, while *Arabidopsis thaliana* exhibited the lowest (∼0.2%) (Figure 1C and S2). Similarly, we observed an average range of 10-15% across all groups for the percentage of Kac proteins, representing the portion of the entire acetylated proteome (Figure 1D and S2). As with Kac sites, *V. cholerae* had the highest percentage (32.7%), followed by the protozoan *Trypanosoma brucei* (24.95%) and the fungus *Aspergillus fumigatus* (22.4%) (Figure S2).

Our observations revealed variations in the percentage of lysine acetylation (Kac) sites and Kac proteins across species. However, it is important to acknowledge limitations. Data normalization attempts to account for proteome size, but differences in methodologies and facilities between laboratories can contribute to observed variations that may not fully reflect biological reality. For example, the *Rattus norvergicus* acetylome identified a higher number of Kac sites (15,474) and Kac proteins (4,541) compared to other species. However, due to its larger proteome, the percentages of total Kac sites and proteins are lower in *R. norvegicus*. Interestingly, notable significant differences persist even within similar taxonomic groups and proteome size ranges (e.g., mammals, bacteria, and fungi). This suggests factors beyond proteome size likely influence Kac abundance.

### Most Kac proteins harbor only a single Kac site

To assess the distribution of lysine acetylation events per protein, we quantified the number of proteins in each acetylome containing 1, 2, 3-5, >5-10, and >10 Kac sites. Combining data across all acetylomes, we found that over 50% of identified Kac proteins have just one detectable Kac site (Figure 2A). The proportion of proteins with 2 or 3-5 Kac sites is around 20%, while those with >5-10 or >10 Kac sites comprise only 6.8% and 1.98%, respectively (Figure 2A).

**Figure 2.**
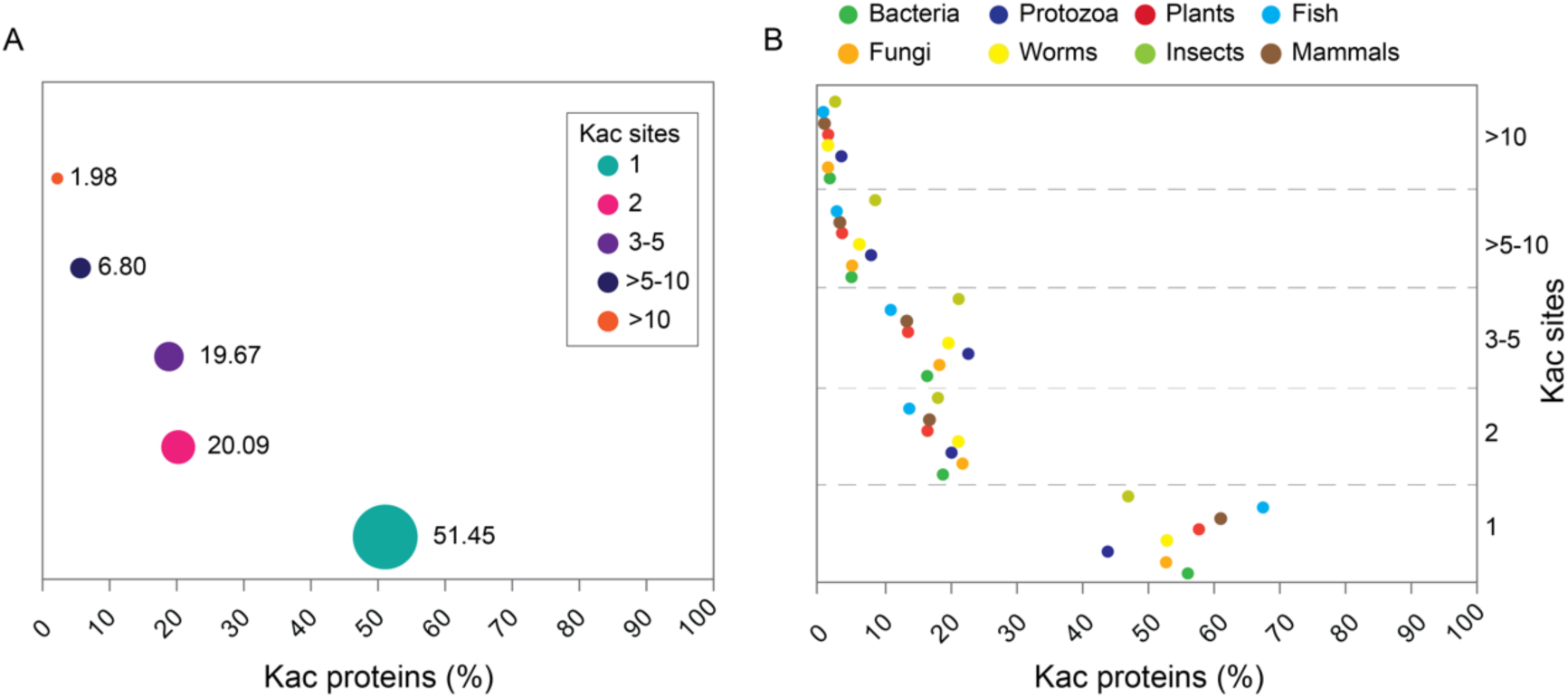
Distribution of Kac sites per protein across the different groups. **A.** Distribution of Kac proteins of all acetylomes analyzed based on the number of Kac sites detected. Most proteins have only one Kac site (50.45 %), while less than 10% of proteins have >5-10 or >10 Kac sites. **B.** Similar results were observed when we analyzed the acetylomes of specific groups (bacteria, fungi, protozoa, plants, worms, fish, insects and mammals) separately.

This trend is also true when analyzing individual organism groups (bacteria, protozoa, plants, fish, fungi, worms, insects, and mammals) (Figure 2B). Fish displayed the highest percentage of proteins with a single Kac site, while protozoa exhibited the highest percentage of proteins with over 10 Kac sites (Figure 2B). Notably, proteins with varying degrees of lysine acetylation (1, 2, 3-5, >5-10, and >10 Kac sites) were identified in all examined species (refer to Figure S3 for detailed information on Kac site distribution per protein in each species). To explore the potential functions of proteins with high lysine acetylation (5, >5-10, and >10 Kac sites), we selected all such proteins from all acetylomes and analyzed their most common protein domains. Notably, those containing the heat shock protein 70 family domain (IPR013126) were present in all three groups (Figure 3). This included 69 proteins from 22 species with 5 Kac sites, 48 proteins from 20 species with >5-10 Kac sites, and 24 proteins from 11 species with over 10 Kac sites. Two additional domain classes were also identified across all groups: transcription factor GTP-binding domain (IPR000795) and transcription elongation factor EFTu-like domain 2 (IPR004161) (Figure 3A). As anticipated, proteins containing histone H2A/H2B/H3 domains were well-represented within the groups harboring 5 or >5-10 Kac sites. These groups accounted for 63 and 58 proteins from 24 species, respectively (Figure 3A).

**Figure 3.**
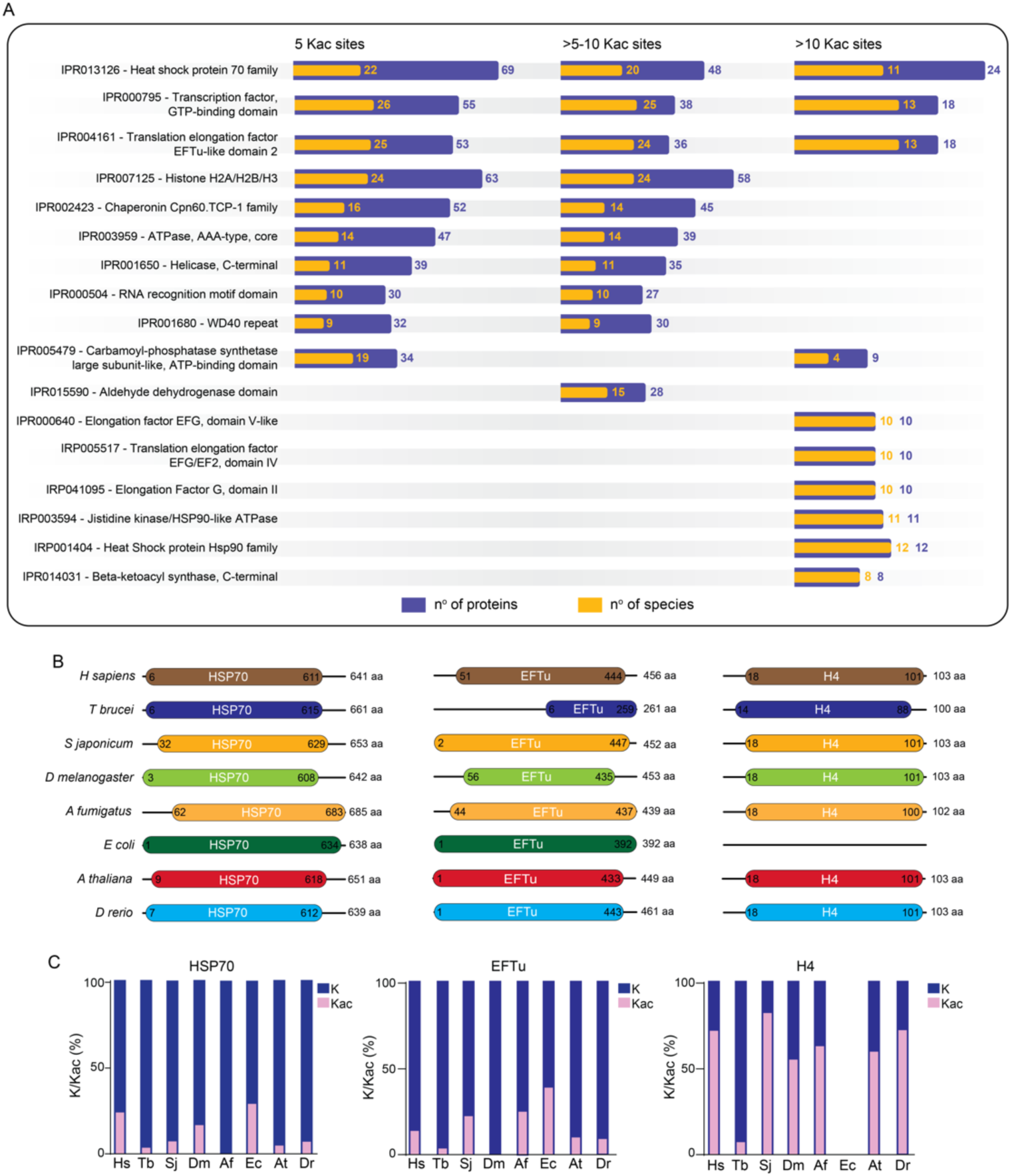
Protein domains associated with high levels of acetylation. **A**. Protein domains enriched with 5 or more Kac Sites. We analyzed the most common protein domains found in proteins identified with 5, >5-10, or >10 Kac sites across all species in the study. Blue bars represent the number of proteins with a specific domain, while orange bars represent the number of species containing that domain within the assessed acetylomes. **B.** Schematic representation of protein size (Hsp70, EFTu, H4) across species. This panel depicts the relative sizes of Hsp70, EFTu, and H4 proteins from different species. **C.** Acetylation levels of Hsp70, EFTu, and H4 proteins across species. Comparison of the percentage of Kac sites in heat shock protein 70 (Hsp70), EFTu, and histone H4 proteins revealed no significant correlation between protein size, total lysine residues, and acetylation level. Blue bars represent the total number of lysine residues, while pink bars represent the percentage of Kac sites found in each protein. Species abbreviations: Hs (*H. sapiens*); Tb (*T. brucei*); Sj (*S. japonicum*); Dm (*D. melanogaster*); Af (*A. fumigatus*); Ec (*E. coli*); At (*A. thaliana*); Dr (*D. rerio*).

We investigated whether the high number of Kac sites observed in Hsp70, EFTu, and H4 proteins correlated with protein size or total lysine residues. We selected representative proteins from various organisms (*H. sapiens*, *T. brucei*, *S. japonicum*, *D. melanogaster*, *A. fumigatus*, *E. coli*, *A. thaliana*, and *D. rerio*) for analysis. Interestingly, while Hsp70 exhibited the highest number of Kac sites and the largest size compared to the medium-sized EFTu and the smaller H4, the overall acetylation percentage of Hsp70 was not as significant as that of H4 (Figure 3B and C). This finding suggests that protein size and total lysine number are not sole determinants of protein acetylation levels.

### Lysine acetylation modifies proteins across various cellular compartments, impacting biological processes and molecular pathways

To gain deeper insights into the role of lysine acetylation in cellular regulation, we combined data from all 48 acetylomes and performed gene enrichment analyses based on Gene Ontology (GO) for cellular components, biological processes, and KEGG pathways.

The cellular component categories with the highest fold enrichment were the proteasome core complex, nucleosome, and mitochondrial F(o) complex (coupling factor for ATP synthase) (Figure 4A). These findings align with the most highly acetylated biological processes, including the ATP metabolic process, tRNA aminoacylation, proton transmembrane transport, proteolysis involved in protein catabolism, and the glycolytic process (Figure 4B). Finally, the most enriched KEGG pathways were related to metabolism, with the top three being the citrate cycle (TCA cycle), propanoate metabolism, and glycolysis (Figure 4C). Details about the enriched GO cellular components, biological processes and KEGG pathways of each specific group of organisms is presented in the Figures S4, S5 and S6.

**Figure 4.**
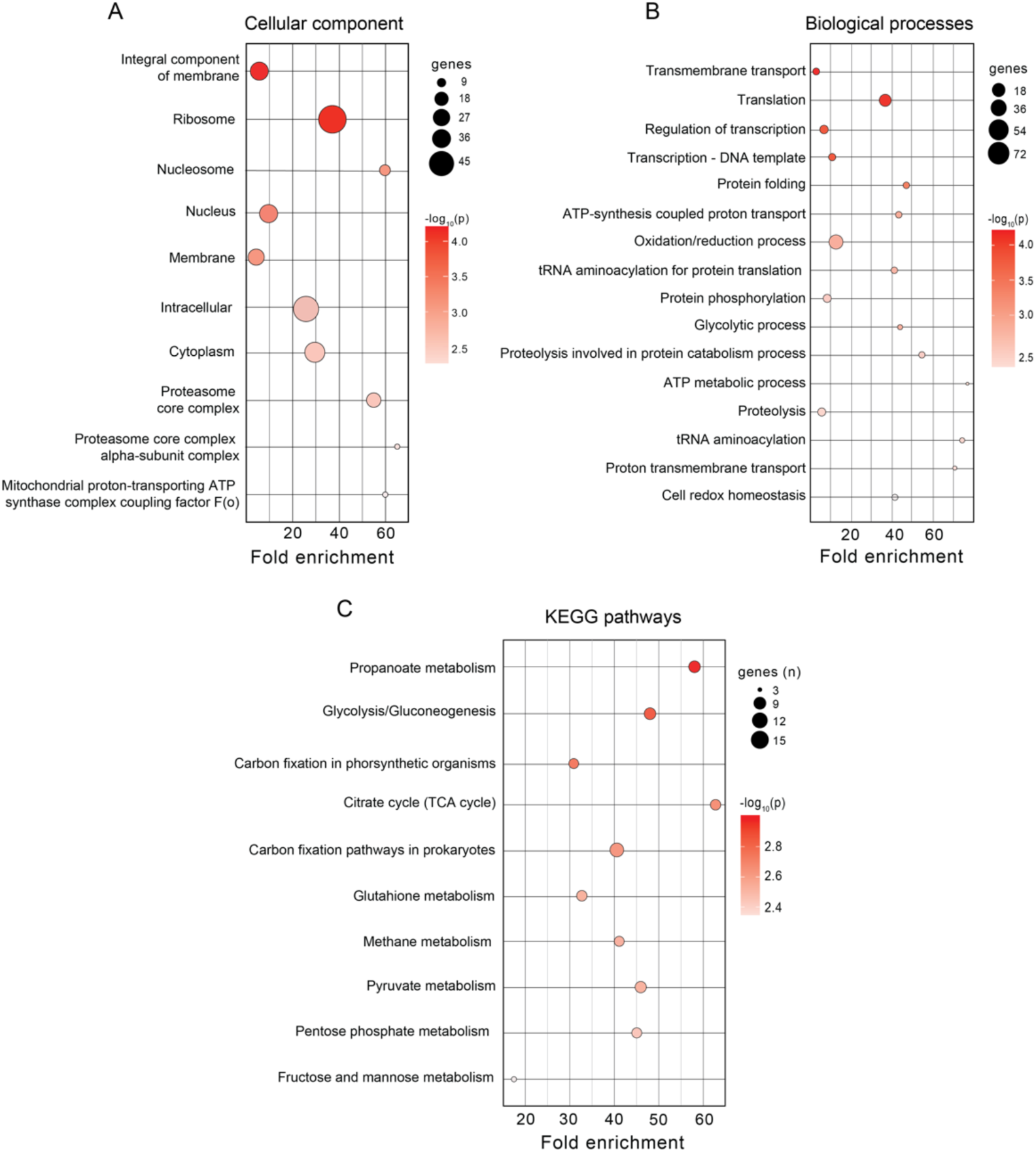
**Gene enrichment analyses of acetylome data. A. Cellular component enrichment**. This panel displays the cellular component categories exhibiting the highest fold enrichment in our gene enrichment analyses. **B. Biological process enrichment**. This panel shows the biological processes most enriched across all studied acetylomes. **C. KEGG pathway enrichment**. This panel depicts the KEGG pathways exhibiting the highest enrichment within the entire group of acetylomes.

These findings suggest a potentially broad regulatory role for protein acetylation. To investigate further this mechanism, we selected some of these cellular processes for in-depth assessment. These processes will be discussed in greater detail in the following sections.

### Heat-shock protein 70 acetylation

Our analysis of domains present enriched in the highly acetylated proteins (containing 5, >5-10, or >10 Kac sites) revealed the presence of heat shock protein 70 (Hsp70) domain-containing proteins in all categories (Figure 3). Hsp70 was originally identified due to its induction by heat stress, and it’s now known to play a crucial role in maintaining cellular homeostasis and promoting cell survival under adverse conditions [16].

To explore the connection between lysine acetylation and Hsp70 proteins, we selected representative species from each group of acetylomes (*T. brucei*, *S. japonicum*, *D. melanogaster*, *E. coli*, *A. thaliana*, *O. sativa*, *S. cerevisiae, A. fumigatus*, and *D. rerio*) for further investigation. As expected, phylogenetic analyses grouped the Hsp70 proteins into distinct clusters (Figure 5A). Amino acid identity comparisons revealed a high degree of sequence similarity among all species, with *D. rerio* (Dr) exhibiting over 80% identity with the human protein (Figure 5B).

**Figure 5.**
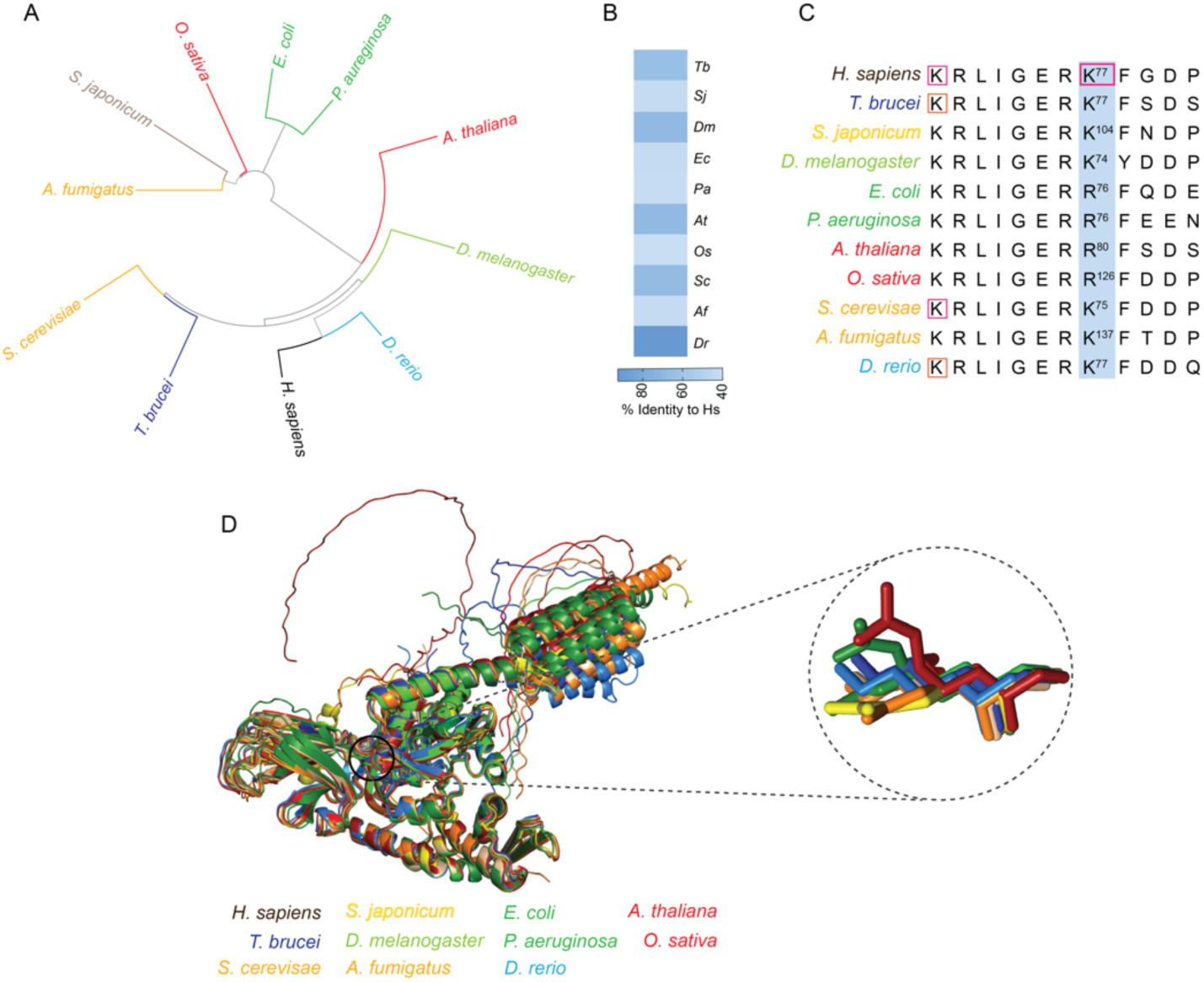
Conservation of the lysine residue regulating Heat Shock Protein 70 activity across species. **A.** Phylogenetic tree of Hsp70 proteins. This panel depicts a phylogenetic tree constructed using heat shock protein 70 (Hsp70) sequences from representative species within each organism group analyzed in this study. **B.** Hsp70 sequence conservation. This panel shows the degree of conservation between the Hsp70 protein sequences of *T. brucei* (Tb), *S. japonicum* (Sj), *D. melanogaster* (Dm), *E. coli* (Ec), *A. thaliana* (At), *O. sativa* (Os), *S. cerevisiae* (Sc), *A. fumigatus* (Af), and *D. rerio* (Dr) compared to the *H. sapiens* protein. **C.** Amino acid alignment of K77 region. This panel displays an amino acid alignment of the K77 residue and flanking regions from the Hsp70 proteins of *H. sapiens* and its orthologs, highlighting the conserved nature of this segment. **D.** Protein structural analysis. This panel presents a comparison of the protein structures for the *H. sapiens* heat shock protein 70 with those of *T. brucei*, *S. japonicum*, *D. melanogaster*, *E. coli*, *A. thaliana*, *O. sativa*, *S. cerevisiae*, *A. fumigatus*, and *D. rerio*.

Interestingly, despite the significant diversity among the chosen organisms, an Hsp70 protein region with high conservation across all species was identified (KRLIGERKFGDP motif, Figure 5D). This region has been shown to be a functionally important regulatory domain in the human protein [17], encompassing the K77 residue, known to be acetylated and involved in the regulation of human Hsp70 (Figure 5C). This residue is conserved in most analyzed species (Figures 5C and D), except for *E. coli*, *P. aeruginosa*, *A. thaliana*, and *O. sativa*, where an arginine replaces the lysine residue (Figure 5C). Importantly, this is a conservative amino acid substitution, since both arginine and lysine are positively charged, sharing structural similarities, which does not rule out the potential regulatory role of that conserved motif for Hsp70 in the different species assessed herein.

### Impact of protein acetylation on the glycolysis

Considering that lysine acetylation has been implicated as a major regulatory cellular mechanism of glucose metabolism [18–21], we decided to determine the glycolytic enzymes identified as acetylated in all acetylomes selected for this study. For this analysis, we used some core glycolytic enzymes: hexokinase (HK), glucose phosphate isomerase (PGI); phosphofructokinase (PFK); fructose-1,6-bisphosphate aldolase (ALD); glyceraldehyde-3-phosphate dehydrogenase (GAPDH), triose-phosphate isomerase (TIM); phosphoglycerate kinase (PGK); phosphoglycerate mutase (PGM); enolase (ENO); pyruvate kinase (PK). Initially, we determined the presence of acetylation in each of the enzymes, considering all the species in each group, and found that at least one glycolytic enzyme is acetylated (an enzyme was considered acetylated regardless the number of Kac sites detected) in at least one of the at least one of the organism’s groups (Figure 6A). We also identified enzymes that were not acetylated, such as PFK and PGK in fish and insects (Figure 6A). Moreover, we identified enzymes within the same group in which acetylation was absent depending on the species, such as TIM in *H. mediterranei* (Figure 6A).

**Figure 6.**
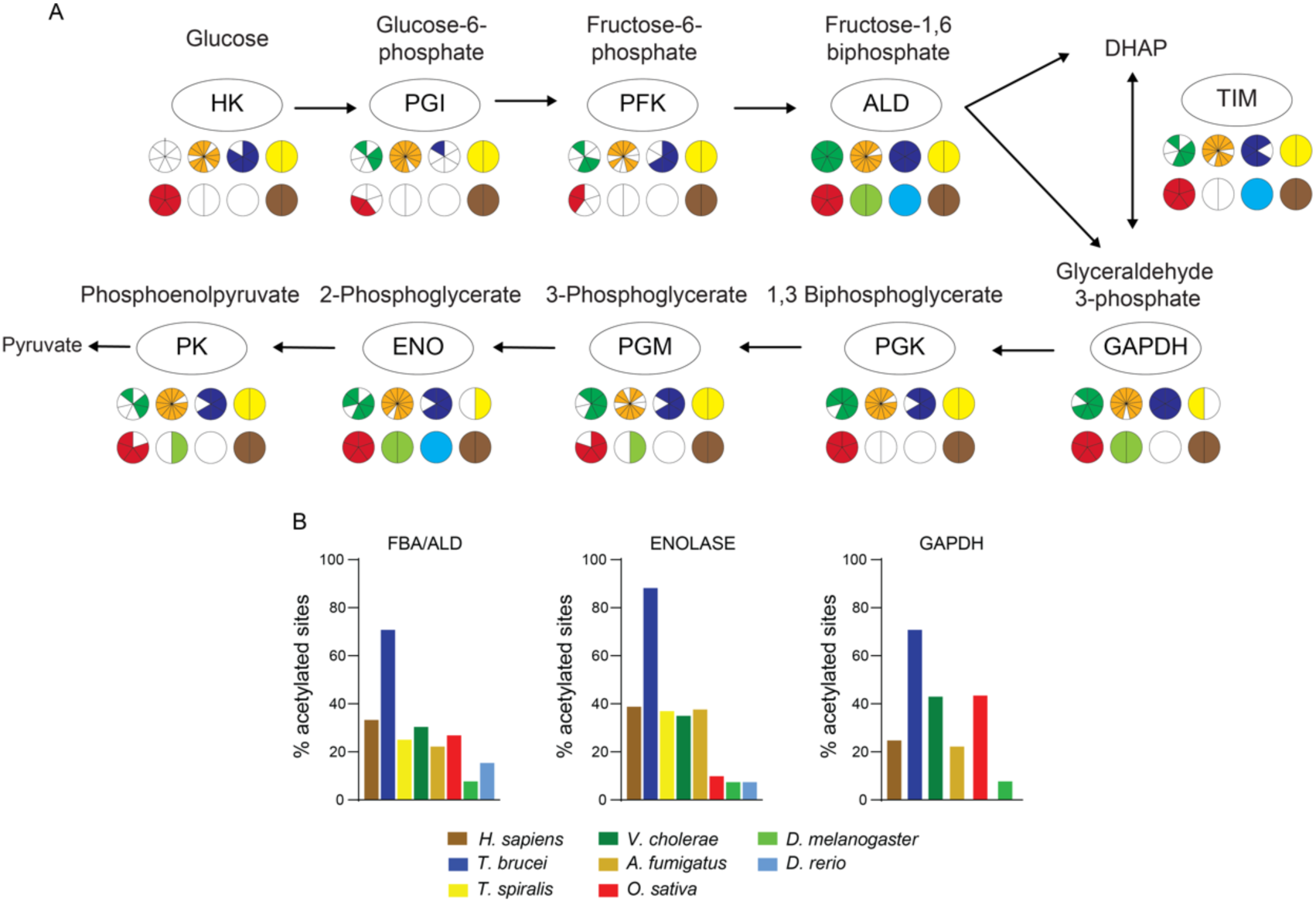
Widespread acetylation of glycolytic enzymes across diverse organisms. **A.** This panel displays the prevalence of acetylation across various glycolytic enzymes within all organism groups analyzed in this study. Each group is represented by a circle below the specific enzyme, with slices within each circle signifying individual species belonging to that group. Color coding is used to differentiate between bacteria (dark green), fungi (orange), protozoa (dark blue), worms (yellow), plants (red), insects (light green), fish (light blue), and mammals (brown). Enzymes were considered acetylated even if only a single Kac site was identified in the specific acetylome. HK (hexokinase), GPI (glucose phosphate isomerase), PFK (phosphofructokinase), ALD (fructose-1,6-bisphosphate aldolase), GAPDH (glyceraldehyde-3-phosphate dehydrogenase), TIM (triose-phosphate isomerase), PGK (phosphoglycerate kinase), PGM (phosphoglycerate mutase), ENO (enolase), PK (pyruvate kinase). **B.** Acetylation levels of representative glycolytic enzymes. This panel presents the percentage of acetylation observed for ALDO, ENO, and GAPDH proteins in representative species from each organism group. Notably, *T. brucei* exhibited the highest level of acetylation for all three enzymes.

A detailed analysis shows that the least acetylated enzymes among the groups were HK, PGI and PFK, while the most acetylated enzyme was ALDO (Figure 11A). Almost no glycolytic enzymes were found acetylated in *D. rerio*, likely due to the low number of only 189 Kac proteins detected in the acetylome of this organism (see Table S1).

To further investigate the acetylation of glycolytic enzymes, we measured Kac levels by determining the percentage of lysine (K) residues acetylated relative to the total number of lysine residues within each enzyme across all species (Figure S7). Figure 6B presents three representative enzymes, enolase (ENO), aldolase (ALDO), and glyceraldehyde-3-phosphate dehydrogenase (GAPDH), which were identified as acetylated in a wider range of species. By selecting a representative species from each group, we observed that the protozoan *T. brucei* exhibited the highest overall acetylation percentage for all three enzymes compared to the other species.

### Lysine acetylation impacts the aldolase catalytic site structure

Fructose-1,6-bisphosphate aldolase (aldolase) is a key glycolytic enzyme. It catalyzes the reversible cleavage of fructose-1,6-bisphosphate into dihydroxyacetone phosphate (DHAP) and glyceraldehyde-3-phosphate (G3P) [22]. The regulatory effect of lysine acetylation on the aldolase activity has been demonstrated for human and *T. brucei* enzymes. It was observed that acetylation at K147 (human enzyme) or K157 (*T. brucei* enzyme) residues abolished the enzyme activity [18, 23], suggesting an evolutionary conserved regulatory mechanism.

To explore the potential regulation of aldolase by acetylation, we selected aldolase enzymes from representative species across various organism groups (*D. melanogaster*, *D. rerio*, *T. brucei*, *T. cruzi*, *A. thaliana*, *T. spiralis*, and *H. sapiens*). We then analyzed their protein structure and acetylation patterns, focusing on the lysine residue known to regulate enzymatic activity in mammals (K147, as described by [18]. Structural comparisons of the predicted enzyme structures from the six species revealed a high degree of conservation in their structures (Figure 7A). Similarly, analysis of both K147 and K230 residues (*H. sapiens*), and surrounding residues involved in enzymatic activity showed that they were conserved (Figure 7B). In this broader analysis, we included sequences from additional species within each organism group.

**Figure 7.**
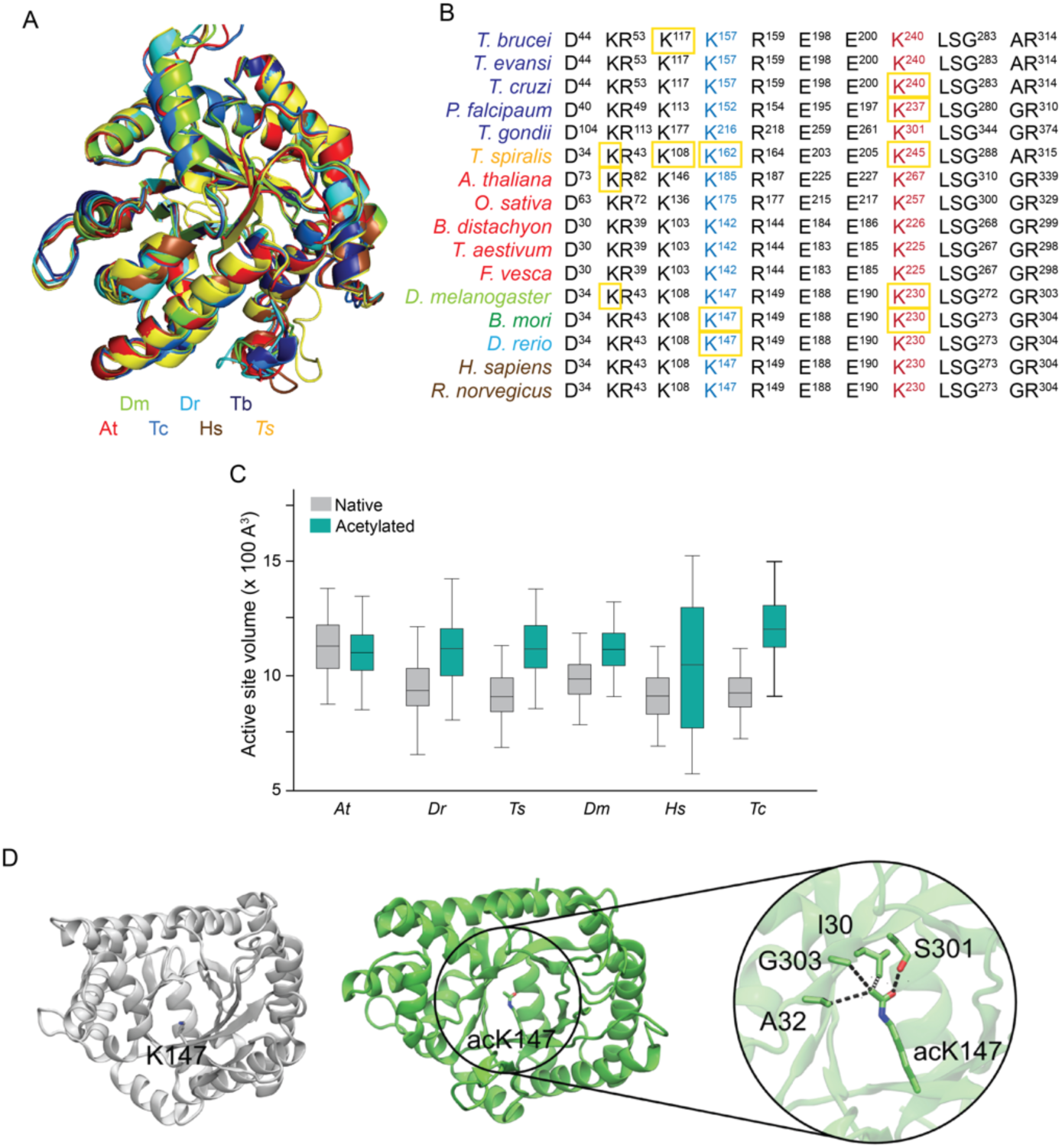
Acetylation of the main lysine residue for aldolase activity affects catalytic site volume and is conserved among different species. **A.** Protein structural comparison of aldose from *A. thaliana* (red), *D. melanogaster* (light green), *D. rerio* (light blue), *T. cruzi* (dark blue) and *H. sapiens* (brown) shows a high degree of structural conservation. **B.** Comparative amino acid alignment of residues belonging to the catalytic site of aldolase from different species. The K147 and K230 residues from *H. sapiens* enzyme involved in the substrate binding are highly conserved among all organisms. Yellow squares indicate the K residues detected acetylated in the corresponding acetylomes. **C.** Effect of acetylation in the aldolase catalytic site volume. The aldolase in its native and acetylated state at the key catalytic lysine residue from *H. sapiens* (K147), *A. thaliana* (K185), *D. rerio* (K147), *T. spiralis* (K162), *D. melanogaster* (K147) and *T. cruzi* (K157), were submitted to molecular dynamic analyses to measure the effect of acetylation on enzyme catalytic site volume. Acetylation increases catalytic site volume in all analyzed species, suggesting a possible impact on enzyme activity as observed for *H. sapiens*. At (*A. thaliana*); Dr (*D. rerio*); Dm (*D. melanogaster*); Hs (*H. sapiens*); Ts (*T. spiralis*) and Tc (*T. cruzi*). **D.** *H. sapiens* aldolase structure, native and K147ac, highlighting the K147 interacting residues involved in substrate binding and enzyme activity.

To gain insight into the regulatory effect of K147 residue acetylation, we performed Molecular Dynamics (MD) simulations to evaluate the impact of this modification in the catalytic site volume, using the native aldolase protein of *D. rerio*, *A. thaliana*, *T. spiralis*, *D. melanogaster*, *H. sapiens* and *T. cruzi*, and its acetylated version at the corresponding K147 residue. Alterations in the catalytic site volume could affect the enzyme-substrate interaction, suggesting an explaination for the inhibitory phenotype observed in human and *T. brucei* aldolase. In general, the acetylated aldolases showed a subtle increase in volume, from ∼950 to 1050 A^3^, except for *A. thaliana*, which showed and average volume of 1129.88 (132,52 SD) and 1104 (121.32 SD) A^3^, respectively (Figure 7C). The largest variation was observed for human aldose, which showed a change in the volume from 924.322 (113.54 SD) (native) to 1060.34 (291.80 SD) (acetylated) (Figure 7C).

Aiming at determining structural aspects related to aldolase acetylation, we measured the distance of the constituent residues to neighbors of the active site (K147) in the human protein. We observed that acetylated K147 (acK147) was projected into a small hydrophobic cavity to interact with I30, A32, and G303 residues, while engaging interactions polar contacts between the carbonyl group in the acetylation and the side chain of the S301 residue (Figure 7D).

Altogether, these results indicate that the post-translational regulatory mechanism of glycolytic enzyme activities might be mainly by altering key catalytic lysine residues, inducing conformational changes in neighbors’ residues and further modulating the enzyme’s function. Also, it is likely that this mechanism has been maintained throughout evolution and potentially plays a role in regulating other glycolytic enzymes activities.

### TCA cycle enzymes are acetylated across species

We also decided to evaluate the presence of acetylation in another crucial pathway for cellular metabolism, the tricarboxylic acid (TCA) cycle, or Krebs cycle. The TCA cycle takes place inside the mitochondria and involves the participation of eight main proteins: citrate synthase (CS), aconitate (ACO), isocitrate dehydrogenase (IDH), α-ketoglutarate (ODH), succinyl-CoA synthetase (SCS), succinate dehydrogenase (SDH), fumarate hydratase (FH) and malate dehydrogenase (MDH) [19, 24]. All the TCA enzymes were found to be acetylated in at least one species of each group, with some variation in the number of species in specific groups (Figure 13). Apart from the group of insects, plants and fish, which had seven and six of the eight enzymes detected acetylated, respectively, the other groups showed acetylation for all eight enzymes (Figure 8). The acetylated proteins found in most species within the groups were CS (acetylated in 28 species from 7 groups), IDH (acetylated in 32 species from 8 groups), and MDH (acetylated in 35 species of 8 groups) (Figure 13 and Sup Table 3). By quantifying the percentage of acetylation of each enzyme per species studied, we found that CS, ACO, SDH, FH and MDH show higher levels of acetylation in mammals; ODH and SCL in bacterial species and bacteria species and IDH in fungi (Figure S5).

**Figure 8.**
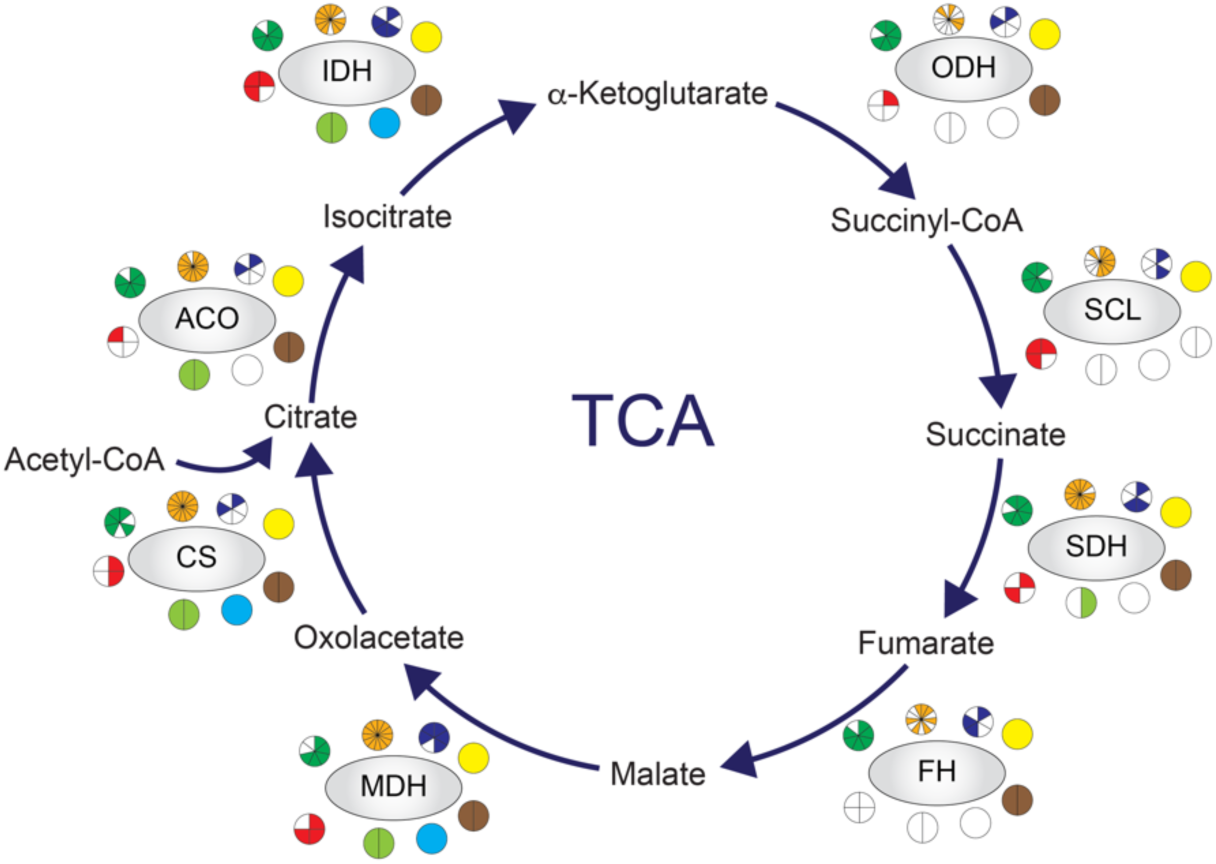
TCA cycle enzymes are highly acetylated in all groups of organisms. Most of the enzymes were detected as acetylated in this study. Each group of organisms is represented by a circle below the specific TCA cycle enzyme and in each circle the slices represent a specie belonging to that group. The groups are represented by different colors as follows: mammals (brown); protozoan (blue); worms (yellow); bacteria (green); fungi (gold); plants (red); insects (light green); fish (cyan). Each enzyme was considered acetylated even if only one Kac site had been identified in the specific acetylome. Citrate synthase (CS), Aconitate (ACO), Isocitrate dehydrogenase (IDH), α-Ketoglutarate (ODH), Succinyl-CoA synthetase (SCL), Succinate dehydrogenase (SDH), Fumarate Hydratase (FH), Malate dehydrogenase (MDH).

### Acetylation in the oxidative stress response

Another cellular process detected in our analysis was oxidation-reduction, represented by different antioxidant enzymes. One such enzyme, superoxide dismutase A (SODA), belongs to a class that detoxifies superoxide into oxygen and hydrogen peroxide. Catalasis ultimately converts hydrogen peroxide into oxygen and water [25].

Previous studies have demonstrated that lysine acetylation regulates SODA activity in human and *T. cruzi* enzymes. Specifically, acetylation of K68 and K97 negatively affects enzyme activity in these organisms, respectively [26, 27]. To investigate whether this regulatory mechanism is conserved throughout evolution, we analyzed SODAs from *A. fumigatus*, *D. melanogaster*, *D. rerio*, *T. cruzi*, *T. brucei*, *A. thaliana*, and *H. sapiens*. We focused on structural conservation and the presence of potential regulatory lysine residues.

Comparative structural alignment of predicted SODA protein structures from these species revealed a high degree of similarity across all enzymes (Figure 9A and Figure S9). Notably, all SODAs possess a conserved “funnel” region that plays a crucial role in directing the substrate towards the enzyme’s active site. This region is typically enriched with positively charged lysine residues, creating a pathway for negatively charged superoxide to reach the catalytic site [26, 27]. Consequently, acetylation of any lysine residue within this region could alter substrate direction due to the neutralization of positive charges [26, 27].

**Figure 9.**
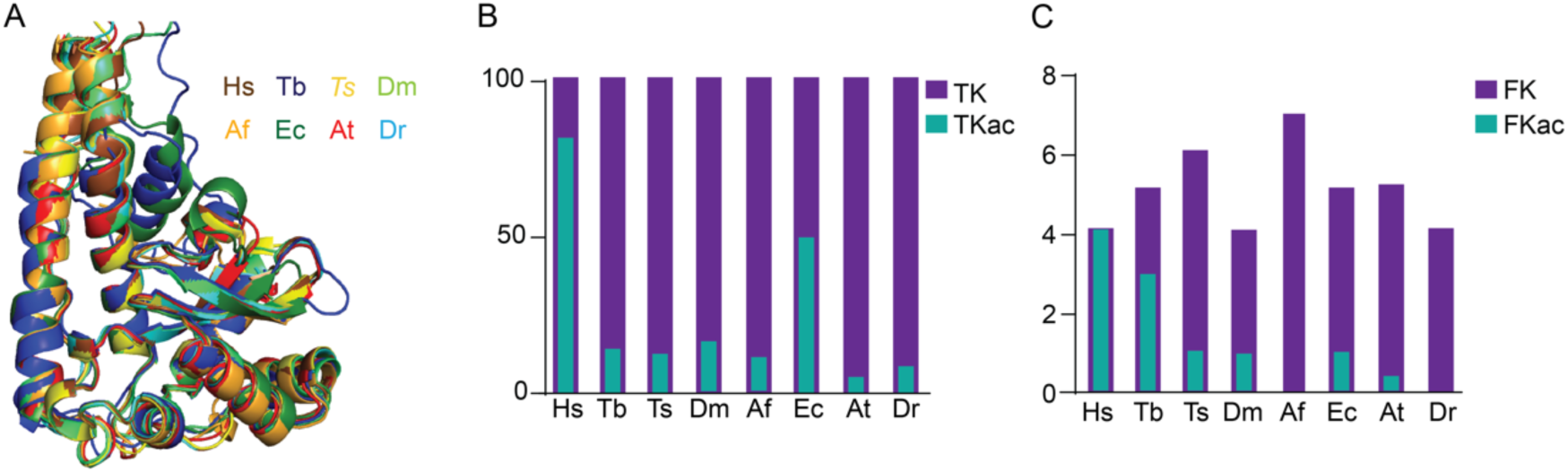
Lysine acetylation of the superoxide dismutase protein in different species. **A.** Comparative analyses of the predicted structures of SODA from *H. sapiens* (Hs); *T. brucei* (Tb); *T. spiralis* (Ts); *D. melanogaster* (Dm); *A. fumigatus* (Af); *E. coli* (Ec); *A. thaliana* (At) and *D. rerio* (Dr), showed a high degree of conservation. **B.** Total percentage of acetylation of each SODA enzyme from the different species. Total lysine residues found in the SODA (TK); Total acetylated-lysine residues found in the SODA. **C.** Number of lysine and acetylated lysine sites present in the regions comprising the funnel that directs the substrate to the enzyme’s catalytic site, suggesting a conserved regulatory mechanism. Lysine residues found acetylated in each SODA (FKac); Total lysine residues (FK).

We quantified the total number of lysine and identified acetylated lysine for each SODA, further determining how many of these acetylated residues were located within the “funnel” region. Human SODA exhibited the highest overall number of acetylated lysines, followed by enzymes from *E. coli*, *D. melanogaster*, and *A. fumigatus* (Figure 9B). Interestingly, when analyzing only the acetylated lysines within the “funnel” region, human SODA again displayed the highest number, but this time followed by *T. brucei* and *D. melanogaster* enzymes (Figure 9C). These findings suggest that, across different species, SODA acetylation within the “funnel” region might play a regulatory role in enzyme activity.

## DISCUSSION

Lysine acetylation, a conserved post-translational modification (PTM) found across diverse organisms, has recently garnered significant interest due to its potential functional roles. Advancements in proteomics and the exploration of metabolic signaling pathways have significantly expanded our understanding of non-histone protein acetylation [6, 7]. This study leverages bioinformatic analysis of published studies, from 2009-2022, describing acetylomes (Figure 1A), highlighting the growing research focus in this field. Notably, a substantial portion of these studies have addressed bacterial acetylation (Figure 1B). The relative ease of studying bacteria in controlled laboratory environments might explain this preference.

This work reveals the significance of acetylation across diverse evolutionary lineages, highlighting notable variability in the proportion of acetylated sites relative to total proteins across several species (Figure S1). Interestingly, bacteria displayed the highest number of acetylation sites, followed by fungi and protozoa (Figure 1C). *Vibrio cholerae*, a bacterium associated with cholera, emerged as the species with the highest acetylation rate (Figure S2). Fungi also displayed a noteworthy percentage of acetylated proteins (Figure 1D). Here it is important to noted that some differences in the acetylation range among the species could also be due to technical differences during the analyses of each work we have selected but could also reflect the physiological states found for each specie in their natural environmental.

Our analysis indicates a relationship between the number of acetylation sites (Kac sites) and their corresponding proteins revealed interesting insights into potential functional implications. Notably, proteins with a single acetylation site dominated across all studied organisms, accounting for approximately 51.45% of the total (Figure 2A). Fish exhibited the highest proportion of single acetylation site detected, followed by insects, plants, and bacteria (Figure 2B). These observations suggest that different organisms might have unique requirements for acetylation, potentially reflecting distinct regulatory mechanisms within their biological processes [7, 8, 28]. Furthermore, the size of proteomes and genomes significantly impacted observed acetylation levels. Mammals, with larger genomes, displayed comparatively lower overall acetylation levels compared to organisms with smaller genomes (Figure S1). Conversely, protozoa, characterized by reduced genomic complexity, exhibited higher acetylation levels, suggesting a potentially compensatory role for acetylation in organisms with smaller genomes.

We further found that the presence of few acetylation sites (2-3) is similar between the groups analyzed (Figure 2B). In addition, proteins from mammalian species presents considerably more acetylation sites, suggesting a potential functional significance of extensive acetylation in mammals, particularly in complex cellular processes and regulatory networks.

HSP70 domains emerged as the most frequently acetylated across species, according to our results (Figure 3). HSP70 proteins are critical for protein folding and unfolding processes, prevention of stress-induced protein aggregation, protein degradation and chaperone-mediated autophagy [29–31]. Evidence accumulated in the last years have shown that Hsp70s are heavily modified at the post-translational level, and that these modifications (phosphorylation, acetylation, methylation, ubiquitination, SUMOylation) fine-tune chaperone function, altering chaperone activity, localization, and selectivity [32–34]. Lysine acetylation was detected in more than 40 sites in Hsp70 from *H. sapiens* and *S. cerevisiae* [35] and our findings demonstrating the strong presence of acetylation, even in species with significant evolutionary and molecular diverge, suggest that acetylation of these domains might directly or indirectly regulate protein activity.

Another finding of our work related to Hsp70 proteins was the conservation of the KRLIGERKFGDP motif (Figure 5D) that bears the K77 residue. The acetylation of K77 by ARD1 acetyltransferase allows Hsp70 to bind to Hop and allowing refolding of denatured proteins [17]. After longer periods of stress, Hsp70 becomes deacetylated, promoting interaction with CHIP to degrade damaged proteins. This switch from protein refolding to degradation is required for the maintenance of protein homoeostasis and protects the cells from stress-induced cell death [17]. We found the K77 present in 7 of 11 species that we analyzed (Figure 5D), suggesting that the regulatory function of acetylation in conserved across the kingdoms. In *E. coli*, *P. aeruginosa*, *A. thaliana*, and *O. sativa* where K77 was replaced by an arginine, that resembles a non-acetylated lysine, it is interesting to speculate that this naturally occurring mutation makes Hsp70 immediately prepared for action to degraded damaged proteins.

As expected, histones, well-known players in epigenetic modifications, were also identified among the most acetylated protein groups (Figure 3). Specifically, H2A, H2B, and H3 histones displayed high levels of acetylation, particularly in proteins with five to ten acetylation sites. However, the detection of histones with more than ten acetylation sites was limited. This might be due to their size or the abundance of lysine and arginine residues, which can be cleaved by trypsin during sample preparation, generating smaller peptides that are difficult to detect by mass spectrometry. The observed abundance of acetylated histones aligns with their established role in post-translational modifications and their crucial function in gene regulation and chromatin remodeling [1, 36, 37].

Our work also indicates the functional significance of lysine acetylation in modulating metabolic processes and molecular pathways. Interestingly, ATP metabolism emerged as the most acetylated process, despite the relatively low number of genes associated with it (Figure 4). This finding highlights the potential importance of acetylation in regulating key cellular functions, as ATP serves as the primary energy source for various processes like signaling, DNA/RNA synthesis, and translation. Similarly, the aminoacyl-tRNA synthetase (aaRS) pathway, crucial for gene expression and protein synthesis, displayed significant acetylation, suggesting a regulatory role for acetylation in this essential process (Figure 4) [19]. Furthermore, oxyreduction, a process intricately linked to glucose degradation and cellular energy generation, exhibited many acetylated genes (Figure 4). This observation further emphasizes the importance of acetylation in regulating core metabolic pathways.

Our analysis focused on metabolic pathways revealed a notable enrichment of acetylation within sugar metabolism pathways, including the tricarboxylic acid (TCA) cycle, propanoate metabolism, and glycolysis (Figure 4B). These pathways are fundamental for energy production and cellular function, highlighting the potential regulatory role of acetylation in metabolic processes[19]. Glucose metabolism has been a major focus in understanding the impact of protein acetylation. Enzymes involved in glucose metabolism, such as those in glycolysis, the TCA cycle, glycogen synthesis, and the irreversible steps of gluconeogenesis, exhibit a high prevalence of acetylation sites [18–20, 23].

Consistent with prior studies, our analysis revealed acetylation of all glycolytic enzymes, with aldolase (ALDO) exhibiting the highest degree of acetylation. Notably, the protein structure and the specific lysine residue (K147 in humans) regulating activity were conserved across analyzed species. This conservation supports the hypothesis that lysine acetylation serves as a general regulator of aldolase activity among diverse evolutionary organisms [18, 23, 28]. These findings suggest that glycolysis is a process regulated by lysine acetylation, regardless of the studied organism’s evolutionary complexity. While further research is necessary, the observed conservation of specific lysine residues within catalytic regions of proteins like aldolase suggests that many other species may also utilize this mechanism for potential enzyme activity regulation [23, 38].

Analysis of the mitochondrial tricarboxylic acid (TCA) cycle, a key cellular process for energy production, also revealed significant levels of acetylation (Figure 8). This cycle comprises eight core proteins, and all were found to be acetylated in at least one group of organisms. Notably, hyperacetylation of the citrate synthase/pyruvate dehydrogenase complex has been shown to negatively regulate the innate immune response in macrophages, thereby impairing the entire TCA cycle function [39]. Furthermore, studies have identified numerous acetylation sites within TCA cycle proteins. These modifications can influence protein activity, with some experiencing increased activity and others experiencing decreased activity [19, 40, 41]. For example, acetylation of *A. thaliana* malate dehydrogenase at K169, K170, and K334 decreases its oxoacetate reduction activity [42], while acetylation of *E. coli* and *H. sapiens* at K99/K140 and K307, respectively, increases its enzymatic activity [43]. These findings highlight the diverse regulatory roles that lysine acetylation can play depending on the specific protein target.

In addition, our findings also revealed the involvement of lysine acetylation in the response to oxidative stress. In particular, the antioxidant enzyme SODA, which has a distinctive funnel-like structure formed by α-helixes that “conduct” the negatively charged superoxide molecule to the catalytic center of the SODA enzyme[44, 45]. The presence of various residues of lysine and arginine at this site propel the superoxide to the catalytic region of the enzyme, carrying out its function of converting superoxide into hydrogen peroxide, reducing cellular damage [25]. Supporting this, previous studies have shown that acetylation at specific lysine residues negatively regulates its activity, such as K68 in humans [27] and K97 in *T. cruzi* [26, 27]. These studies indicate that acetylation alters the positive charge of lysines, which hinders the “conduction” of the superoxide to the catalytic center of the SODA enzyme [26, 27].

Our combined analysis of acetylome data and relevant literature provides strong evidence for the widespread occurrence of lysine acetylation across diverse evolutionary groups. This post-translational modification extends its functional regulatory aspect to critical cellular processes, metabolic pathways, and conserved protein domains. Notably, our findings highlight the importance of protein acetylation in regulating both glucose metabolism, TCA cycle and the oxidative stress response. Several studies indicate that acetylation of key lysine residues in enzymes as aldolase (glycolysis) and superoxide dismutase (antioxidant defense) can play a crucial role in cellular metabolic processes and redox balance [8, 18, 19, 21, 23].

Despite our extensive review and meta-analysis of acetylomes, it is important to bear in mind that the transient nature of acetylation makes it difficult to precisely map acetylation profiles. Therefore, those species in which no protein acetylation was identified are encouraged to be further investigated with more sophisticated techniques [46]. Furthermore, extensive biochemical work is required to validate the role of acetylation in regulating the activity of specific enzymes and its consequently enzymes and consequently cellular processes.

## MATERIALS AND METHODS

### Data Collection and curation

To identify published acetylome data, we conducted a keyword search in PubMed (www.pubmed.ncbi.nlm.nih.gov) using the term “acetylome” and limited the search to publications from 2009 to 2022. This timeframe reflects the appearance of the first published acetylome in 2009. We downloaded all retrieved articles and their supplementary materials containing lists of identified acetylated proteins for further curation and analysis. From this initial search, we identified 128 acetylomes encompassing a broad range of species, including bacteria, fungi, protozoa, worms, plants, insects, crustaceans, fish, and mammals.

To minimize potential variations within our analyses, we subsequently filtered the collected acetylomes. This involved selecting only studies that employed lysine-acetylated peptide enrichment techniques. Additionally, we excluded bacterial data from[15] due to its extensive focus on 48 phylogenetically diverse bacteria. Instead, we opted to include data from other bacterial species within our study. Following these refinements, our final dataset comprised 48 acetylomes representing 37 species across 8 phylogenetic clades.

### Data Organization and Initial Analysis

The collected data underwent initial organization to determine the number of acetylated proteins (Kac proteins) and the number of lysine-acetylated sites (Kac sites). Additionally, we retrieved proteome size information for all analyzed species from either UniProt (https://www.uniprot.org) or VEuPathDB (https://veupathdb.org/veupathdb/app). To minimize bias arising from experimental design variations across different acetylomes, we normalized the Kac sites and Kac proteins identified. This normalization considered the total number of lysine residues within each predicted proteome and the overall proteome size (number of proteins) of each species.

We further categorized the Kac proteins based on the number of identified Kac sites within each acetylome. This manual categorization grouped proteins based on the following criteria 1, 2, 3-5, 5, >5-10 and >10 Kac sites. For each species, the data was compiled into a table, including the UniProt ID for each protein within each category (1, 2, 3-5, 5, >5-10 and >10 Kac sites). The data was then visualized using GraphPrism 9.5, and final figures were generated with Adobe Illustrator.

### Systematic assessment of Kac sites and Kac-containing proteins’ functional enrichment analyses

*Ad-hoc* bash commands were designed for counting sites and retrieving IDs, in a systematic fashion, of all Kac-containing proteins/genes from supplementary tables of articles reporting acetylomes from all organisms studied herein (Table S1). GO-basic.obo and KEGG “htext” files were downloaded from http://current.geneontology.org/ontology/go-basic.obo and https://www.genome.jp/kegg-bin/get_htext?ko00001, respectively. PERL scripts were written to automatically prepare 2x2 contingency tables (https://github.com/eltonjrv/bioinfo.scripts/blob/master/GOcount4fisher.pl, https://github.com/eltonjrv/bioinfo.scripts/blob/master/KOcount4fisher.pl, https://github.com/eltonjrv/bioinfo.scripts/blob/master/assign-GO2VGs.pl) to be used as input for the gene enrichment analysis run in R through a Fisher Exact Test execution (https://github.com/eltonjrv/bioinfo.scripts/blob/master/fisher4GOenrichment.R). Significant GO terms and KEGG biochemical pathways *(p* < 0.01) were plotted on either bar or bubble charts through ggplot2 or pathfinder (Ulgen 2019 - PMID: 31608109) R packages. R environment version 4.1 was used for such purpose.

### Protein structural analyses

In the structural comparison assays, we initially obtained the protein structures using the PDB data bank (https://www.rcsb.org), and for those proteins with no data available, the predicted protein structure was generated using the AlphaFold AI tool [47]. The analyses were done using the software PyMOL and the images generated were processed in Adobe Illustrator.

### Molecular dynamic (MD) simulations of aldolase

The initial three-dimensional structures for each aldolase studied were generated by molecular modeling (*A. thaliana*, 1ADO; *D. rerio*, 1ADO; *T. spiralis*, 1FDJ; and *D. melanogaster*, 1FBA) using SWISSMODEL [48] or obtained by structures generated experimentally (*H. sapiens*, 5KY6; and *T. cruzi*, 1F2J). For each of them, the native and acetylated version was generated to the corresponding amino acid residue 147 of human aldolase.

Twelve structures were generated, where each one was submitted to Molecular Dynamics (MD) simulations, using the GROMACS software [49] under the field of force Charmm36m [50]. The protonation state of each protein was measured for the pH range equal to 7 using the PROPKA server, while the acetylation was performed using the PyTM plugin [51]. Then, each structure was placed in a dodecahedral box 12 Å from the edge of the box to the farthest atom on the XYZ axes. Each system was then solvated, neutralized and equilibrated with 0.1 M NaCl. A minimization step was performed to reduce the system’s energy below 1000 kJ/mol/nm^2^, using the Steepest Descent algorithm. Next, an equilibration step was performed, controlling the temperature at 300 K using the V-Rescale thermostat [52] for 1 ns, then another step of also 1 ns was performed to monitore the pressure with the Berendsen barostat [53] at 1 bar. In the final step, the thermostat and barostat were changed to Nose-Hoover [54, 55]. Three independent replicas of 300 ns each were simulated, collecting frames every 20 ps.

Structural measurements were employed to analyze the effect of acetylation on each protein studied. The active site volume and distance from key residues were measured using Epock software [56] and scripts implemented in VMD [57], respectively. In the first, an 8 Å sphere was positioned over the site cavity, to cover the catalytic site residues, including the acetylated lysine of interest, measuring the volume of each frame collected during MD simulations. In the last one, the minimum distance between the residues was measured, using human aldolase as a reference, D34, K108, K147, R149, E188, E190, K230 and S301.

## Author contribution

N.S.M. conceptualized and acquired funds for the project. BSB designed and performed most of the analyses, analyzed the data, and drafted the manuscript. ABL, SRM, ACCNS contributed to the initial analyses of the acetylomes. E.J.R.V. performed the bioinformatics analyses and revised the manuscript. AASG performed the molecular dynamics analyses and revised the manuscript. N.S.M. supervised the project, wrote, and revised the final version of the manuscript. All authors read and approved the final version

---

**Figure S1.**
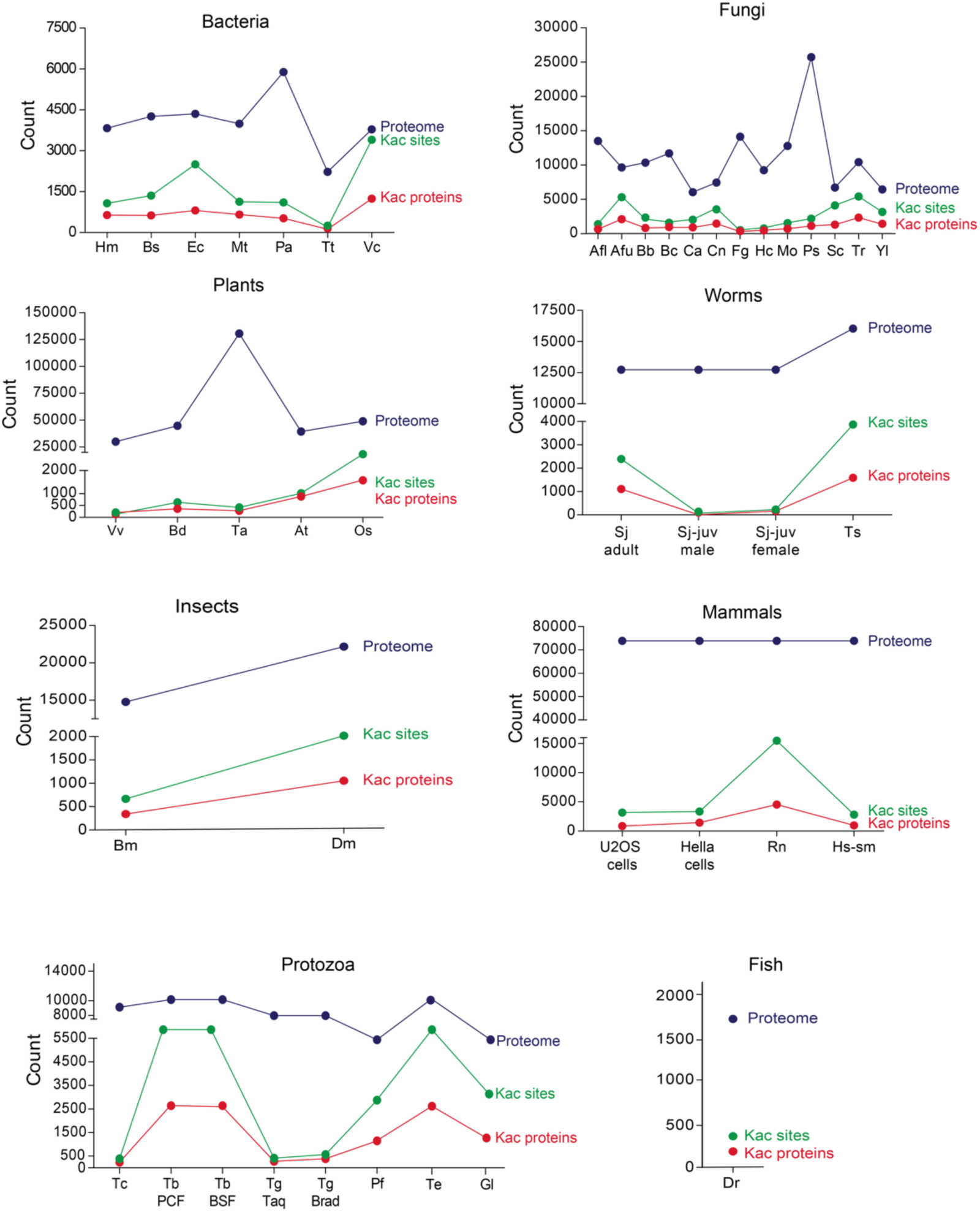
Distribution of acetylomes selected for this work based on proteome size, number of acetylated sizes and acetylated proteins. The acetylome descriptions of all species for each group of organisms (bacteria, fungi, plants, worms, insects, mammals and protozoa) were plotted to give a broad idea about the characteristics of each acetylome. Each plot contains the size of the proteome, the number of acetylated-lysine sites (Kac sites) and acetylated proteins (Kac proteins). Hm (*Haloferax mediterranei*); Bs (*Bacillus subtilis*); Ec (*Escherichia coli*); Mt (*Mycobacterium tuberculosis*); Pa (*Pseudomonas aeruginosa*); Tt (*Thermus thermophilus*); Vc (*Vibrio colerae*); Afl (*Aspergillus flavus*); Afu (*Aspergillus fumigatus*); Bb (*Beauveria bassiana*); Bc (*Botrytis cinerea*); Ca (*Candida albicans*); Cn (*Cryptococcus neoformans*); Fg (*Fusarium graminearium*); Hc (*Histoplasma capsulatum*); Mo (*Magnaporthe oryzae*); Ps (*Phytophthora sojae*); Sc (*Saccharomyces cerevisiae*); Tr (*Trichophyton rubrum*); Yl (*Yarrowia lipolytica*); At (*Arabdopsis thaliana*); Os (*Oryza sativa*); Ta (*Triticum aestivum*); Bd (*Brachypodium distachyon*); Vv (*Vitis vinifera*); Sj adult (*Schistosoma japonicum* adult); Sj-juv male (*Schistosoma japonicum* juvenile male form); Sj-juv female (*Schistosoma japonicum* juvenile female form); Ts (*Trichinella spiralis*); Bm (*Bombyx mori*); Dm (*Drosophila melanogaster*); Tc (*Trypanosoma cruzi*); Tg Taq (*Toxoplasma gondii, tachyzoite form*); Tg Brad (*Toxoplasma gondii, bradyzoite form*); Tb PCF (*Trypanosoma brucei* procyclic form); Tb BSD (*Trypanosoma brucei* bloodstream form); Pf (*Plasmodium falciparum*); Te (*Trypanosoma evansi*); Gl (*Giardia lamblia*); Dr (*Danio rerio*); Rn (*Rattus novergicus*); Hs-sm (*Homo sapiens* skeletal muscle).

**Figure S2.**
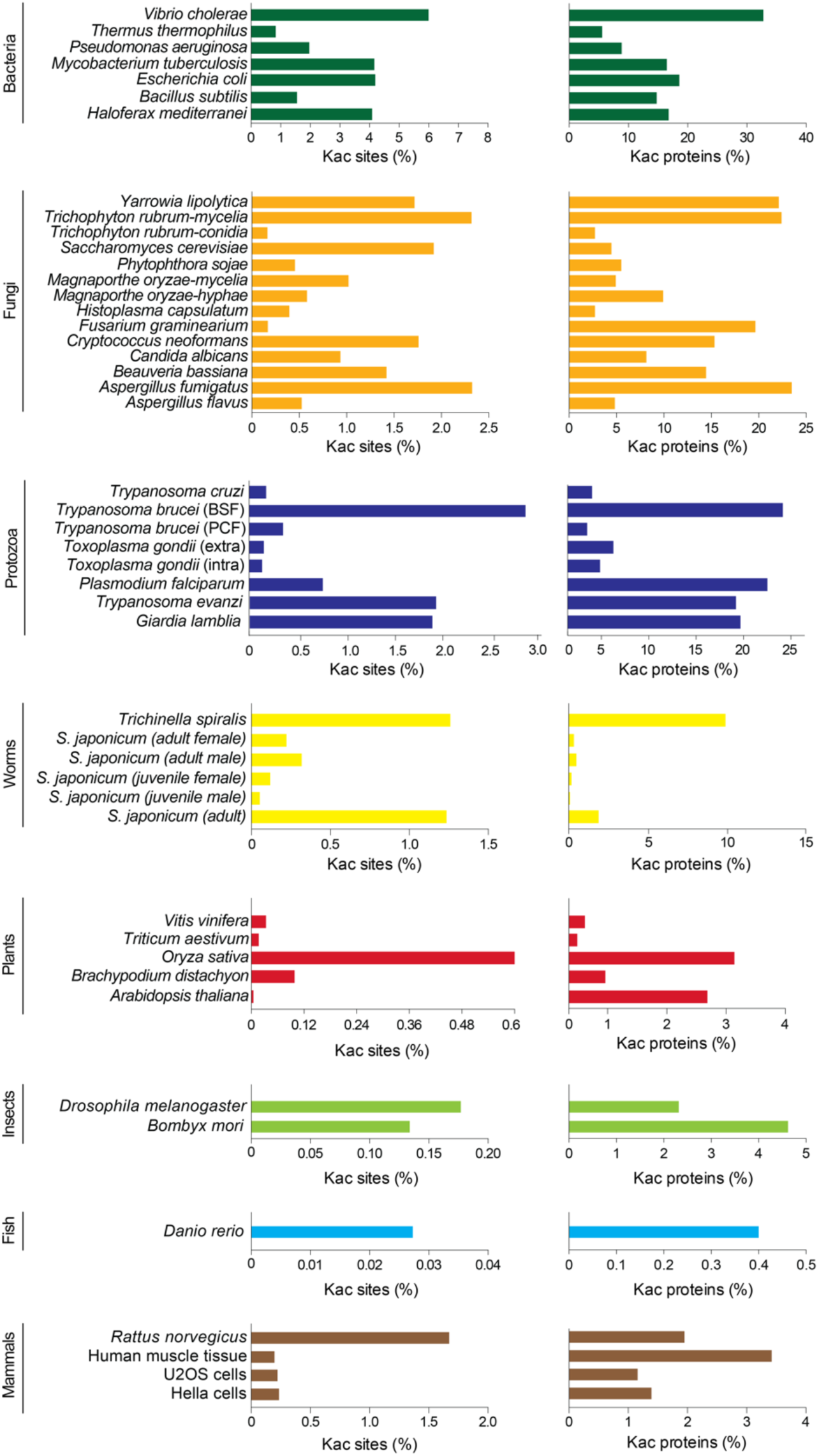
Percentage of the lysine-acetylated sites and lysine-acetylated proteins found in each group of organisms. The percentages of the lysine-acetylated sites and lysine-acetylated proteins were calculated based on the total number of lysine residues found in each proteome and the size of the proteome from each specie.

**Figure S3.**
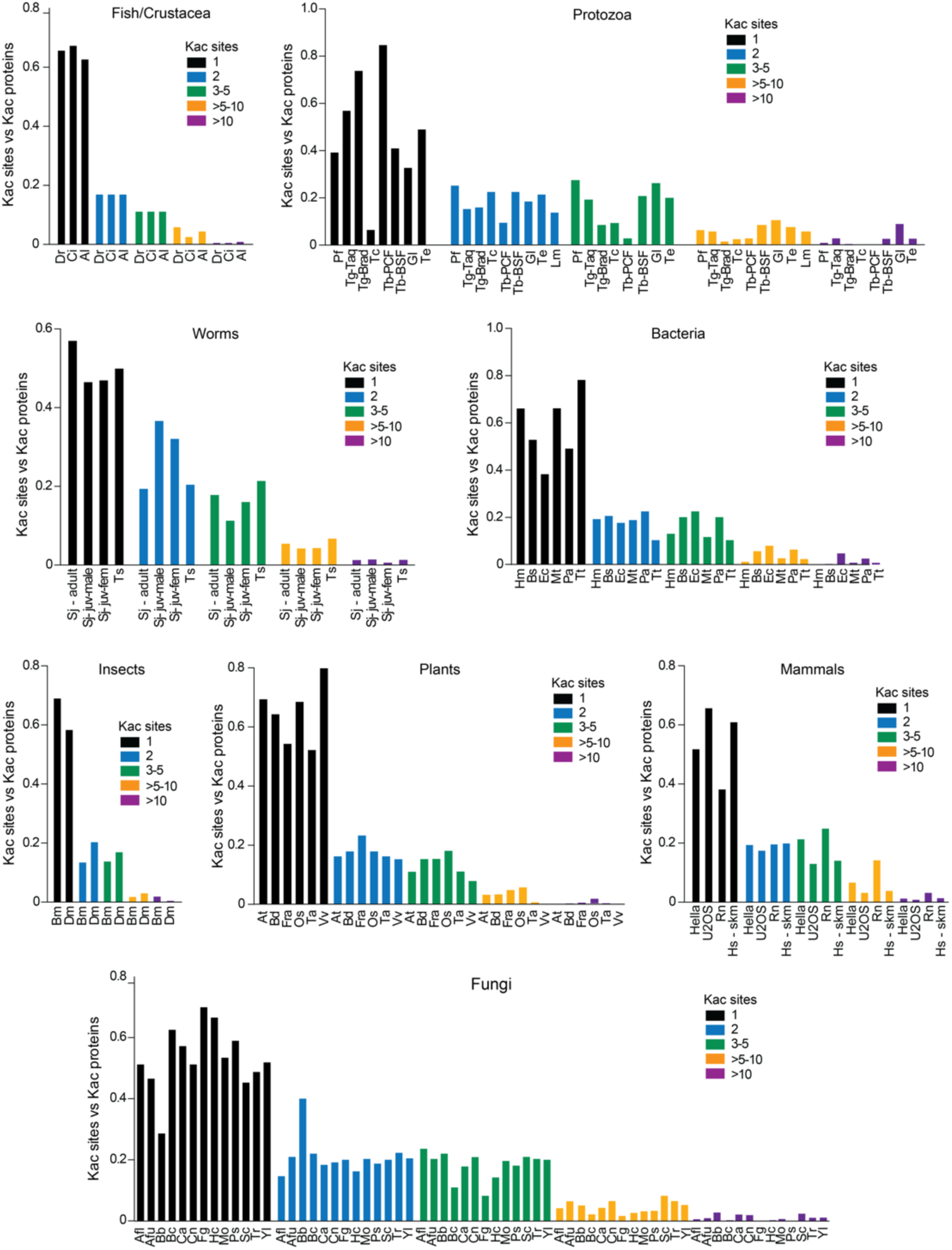
Number of Kac sites detected in each protein in all acetylomes analyzed. The number of proteins identified with 1, 2, 3-5, >5-10 and >10 Kac were quantified using the acetylome data for each specie analyzed. Hm (*Haloferax mediterranei*); Bs (*Bacillus subtilis*); Ec (*Escherichia coli*); Mt (*Mycobacterium tuberculosis*); Pa (*Pseudomonas aeruginosa*); Tt (*Thermus thermophilus*); Vc (*Vibrio colerae*); Afl (*Aspergillus flavus*); Afu (*Aspergillus fumigatus*); Bb (*Beauveria bassiana*); Bc (*Botrytis cinerea*); Ca (*Candida albicans*); Cn (*Cryptococcus neoformans*); Fg (*Fusarium graminearium*); Hc (*Histoplasma capsulatum*); Mo (*Magnaporthe oryzae*); Ps (*Phytophthora sojae*); Sc (*Saccharomyces cerevisiae*); Tr (*Trichophyton rubrum*); Yl (*Yarrowia lipolytica*); At (*Arabdopsis thaliana*); Os (*Oryza sativa*); Ta (*Triticum aestivum*); Bd (*Brachypodium distachyon*); Vv (*Vitis vinifera*); Sj adult (*Schistosoma japonicum* adult); Sj-juv male (*Schistosoma japonicum* juvenile male form); Sj-juv female (*Schistosoma japonicum* juvenile female form); Ts (*Trichinella spiralis*); Bm (*Bombyx mori*); Dm (*Drosophila melanogaster*); Tc (*Trypanosoma cruzi*); Tg Taq (*Toxoplasma gondii, tachyzoite form*); Tg Brad (*Toxoplasma gondii, bradyzoite form*); Tb PCF (*Trypanosoma brucei* procyclic form); Tb BSD (*Trypanosoma brucei* bloodstream form); Pf (*Plasmodium falciparum*); Te (*Trypanosoma evansi*); Gl (*Giardia lamblia*); Dr (*Danio rerio*); Rn (*Rattus novergicus*); Hs-sm (*Homo sapiens* skeletal muscle).

**Figure S4.**
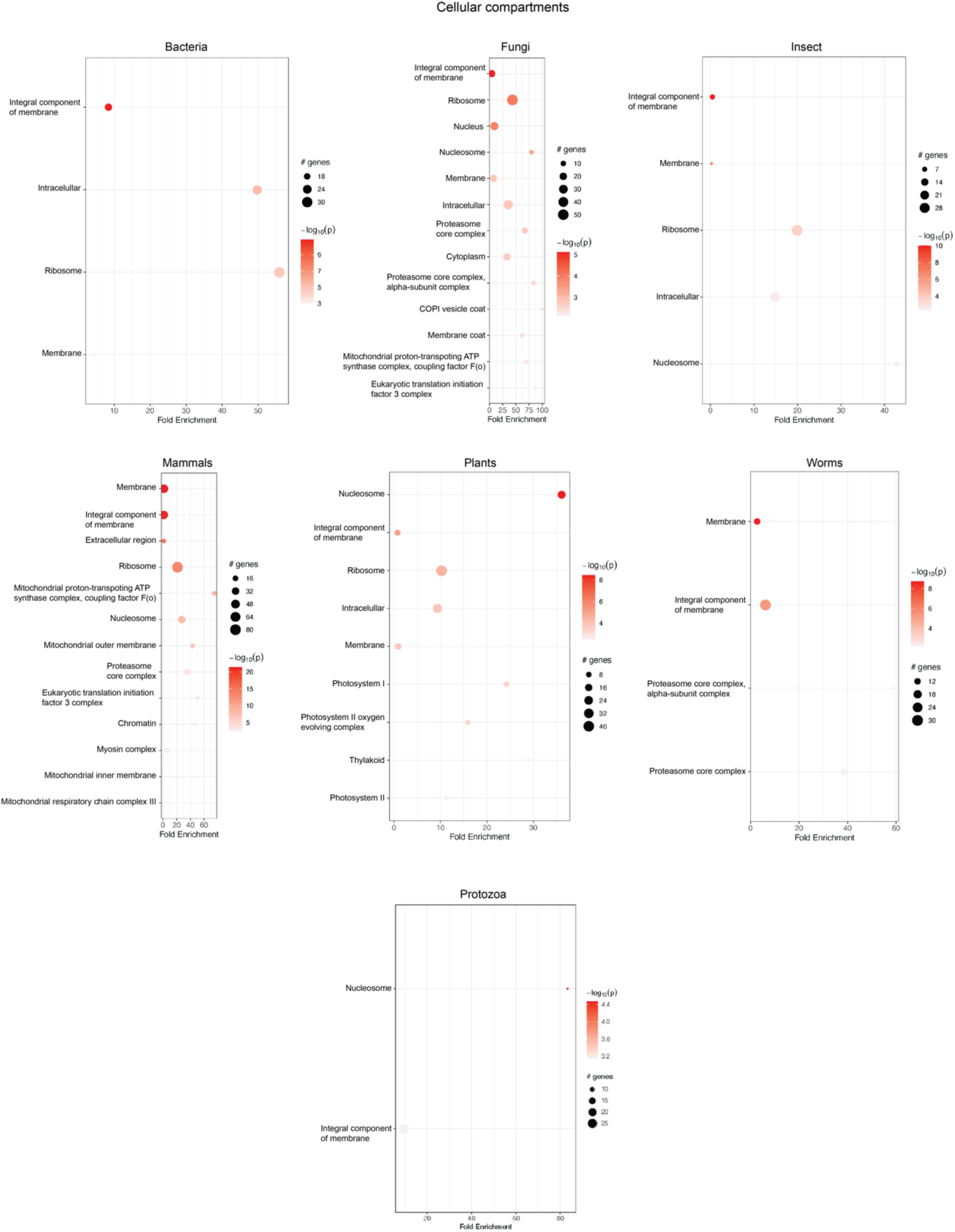
Cellular component enrichment across analyzed groups. Here we demonstrated the cellular component categories enriched in at least two different species within each analyzed group.

**Figure S5.**
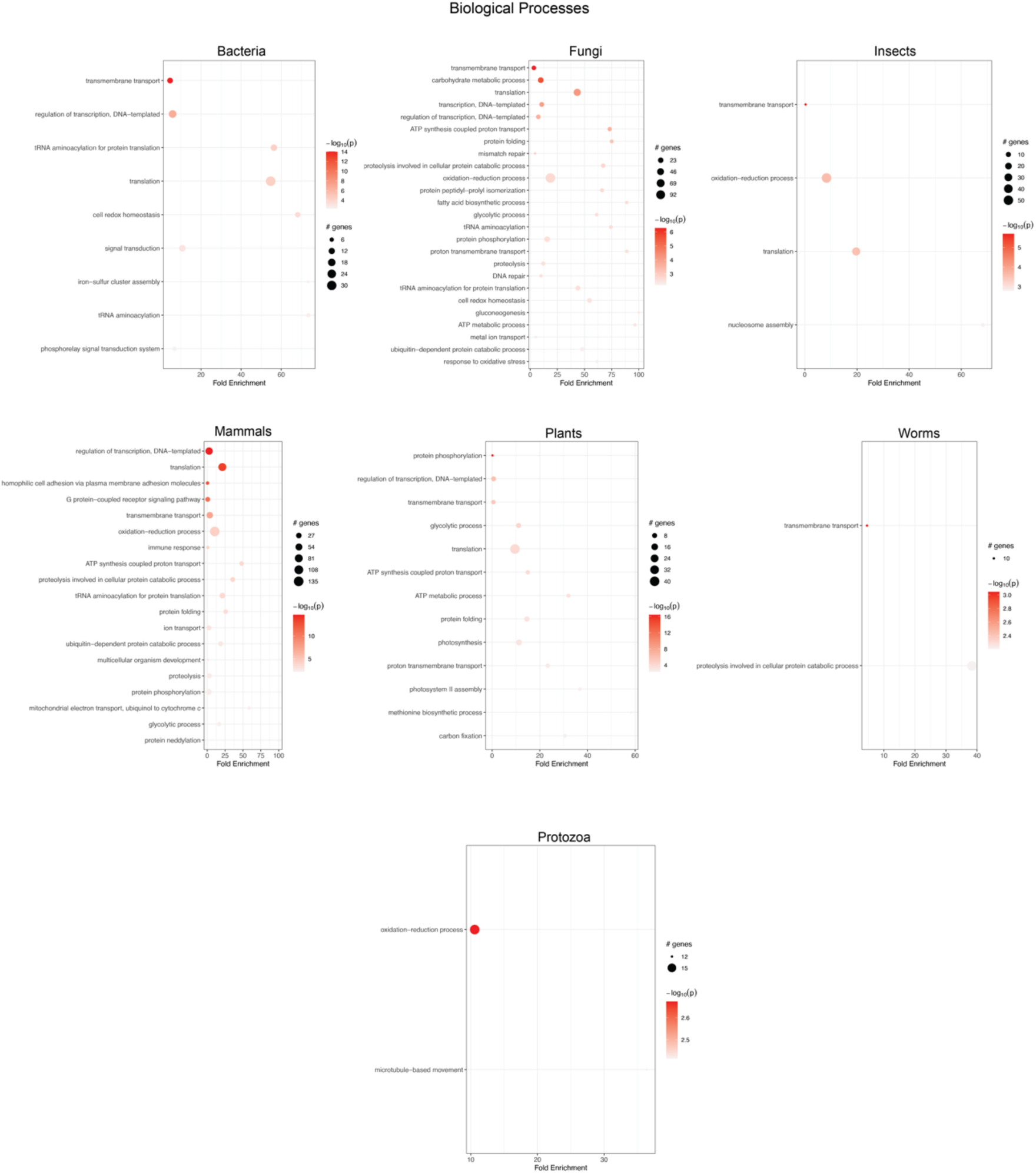
Biological processes enrichment across analyzed groups. Here we demonstrated the biological processes categories enriched in at least two different species within each analyzed group.

**Figure S6.**
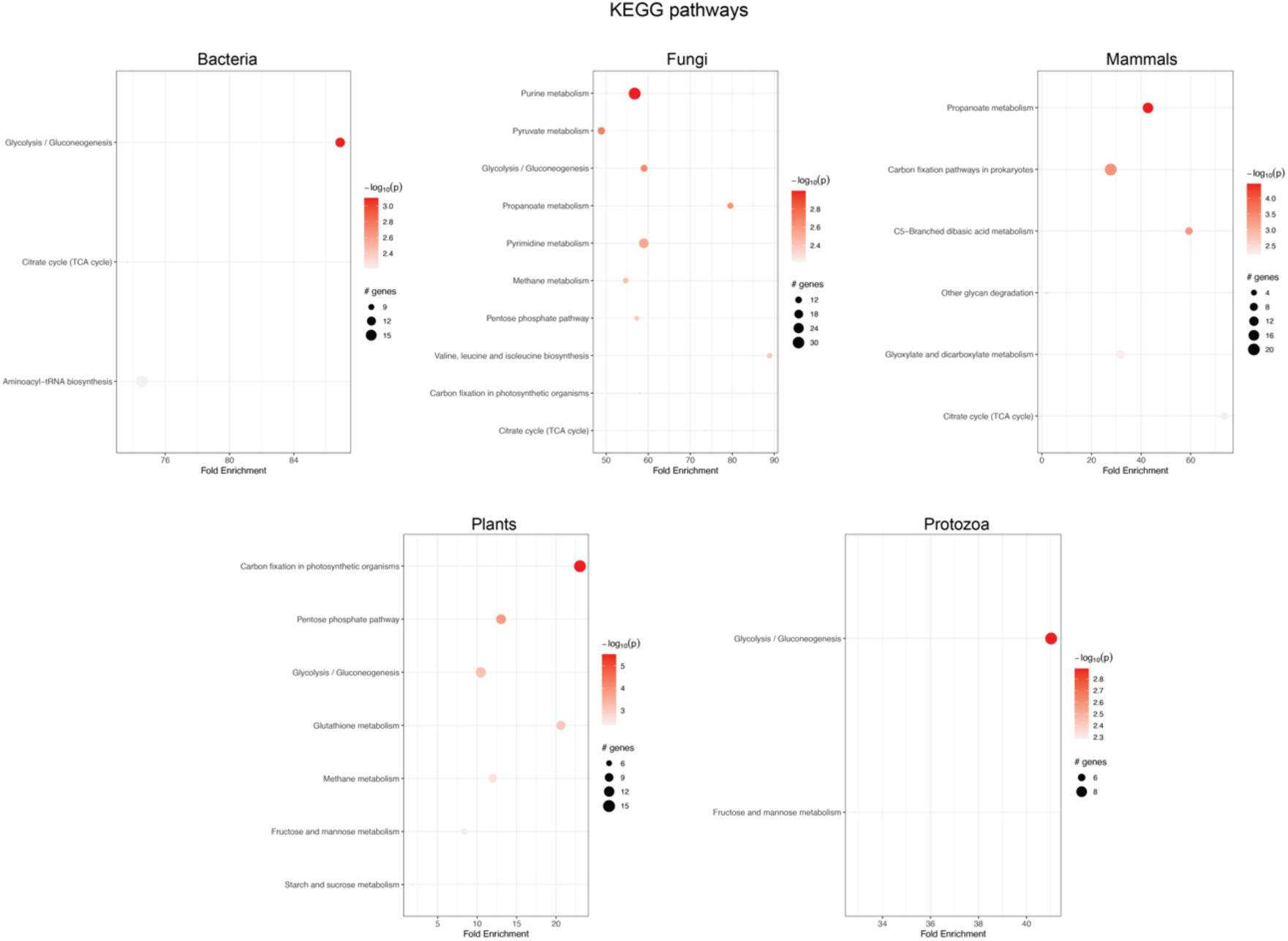
Biological processes enrichment across analyzed groups. Here we demonstrated the KEGG categories enriched in at least two different species within each analyzed group.

**Figure S7.**
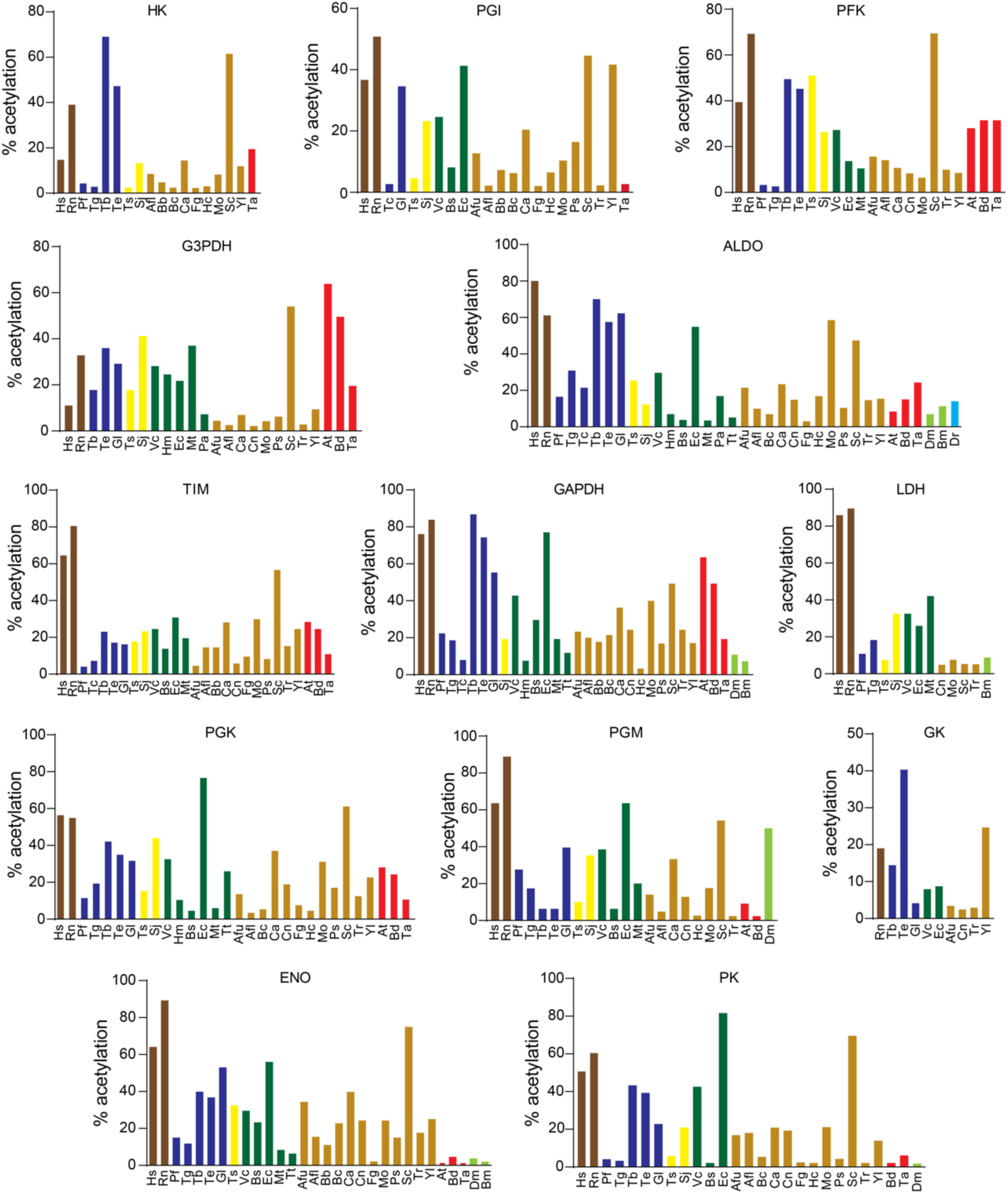
Glycolytic enzymes percentage of the lysine-acetylated and non-acetylated found in each group of organisms of different species. Enzymes were considered acetylated even if only one Kac site had been identified in the specific acetylome. Hexokinase (HK), glucose phosphate isomerase (PGI); phosphofructokinase (PFK); fructose-1,6-bisphosphate aldolase (ALD); glyceraldehyde-3-phosphate dehydrogenase (GAPDH), triose-phosphate isomerase (TIM); phosphoglycerate kinase (PGK); phosphoglycerate mutase (PGM); enolase (ENO); pyruvate kinase (PK). The groups are represented by different colors as follows: mammals (brown); protozoan (blue); worms (yellow); bacteria (green); fungi (gold); plants (red); insects (light green); fish (cyan). Identified species: Hs (*Homo sapiens)*; Rn (*Rattus* novergicus); Tb (*trypanosoma* brucei); Tc (*Trypanosoma cruzi*); Tg (*Toxoplasma gondii*); Pf (*Plasmodium falciparum*); Te (*Trypanosoma evansi*); Gl (*Giardia lamblia*); Hm (*Haloferax mediterranei*); Bs (*Bacillus subtilis*); Ec (*Escherichia coli*); Mt (*Mycobacterium tuberculosis*); Pa (*Pseudomonas aeruginosa*); Tt (*Thermus thermophilus*); Vc (*Vibrio colerae*); Afl (*Aspergillus flavus*); Afu (*Aspergillus fumigatus*); Bb (*Beauveria bassiana*); Bc (*Botrytis cinerea*); Ca (*Candida albicans*); Cn (*Cryptococcus neoformans*); Fg (*Fusarium graminearium*); Hc (*Histoplasma capsulatum*); Mo (*Magnaporthe oryzae*); Ps (*Phytophthora sojae*); Sc (*Saccharomyces cerevisiae*); Tr (*Trichophyton rubrum*); Yl (*Yarrowia lipolytica*); At (*Arabdopsis thaliana*); Os (*Oryza sativa*); Ta (*Triticum aestivum*); Bd (*Brachypodium distachyon*); Vv (*Vitis vinifera*); Sj (*Schistosoma japonicum*); Ts (*Trichinella spiralis*); Bm (*Bombyx mori*); Dm (*Drosophila melanogaster*); Dr (*Danio rerio*).

**Figure S8.**
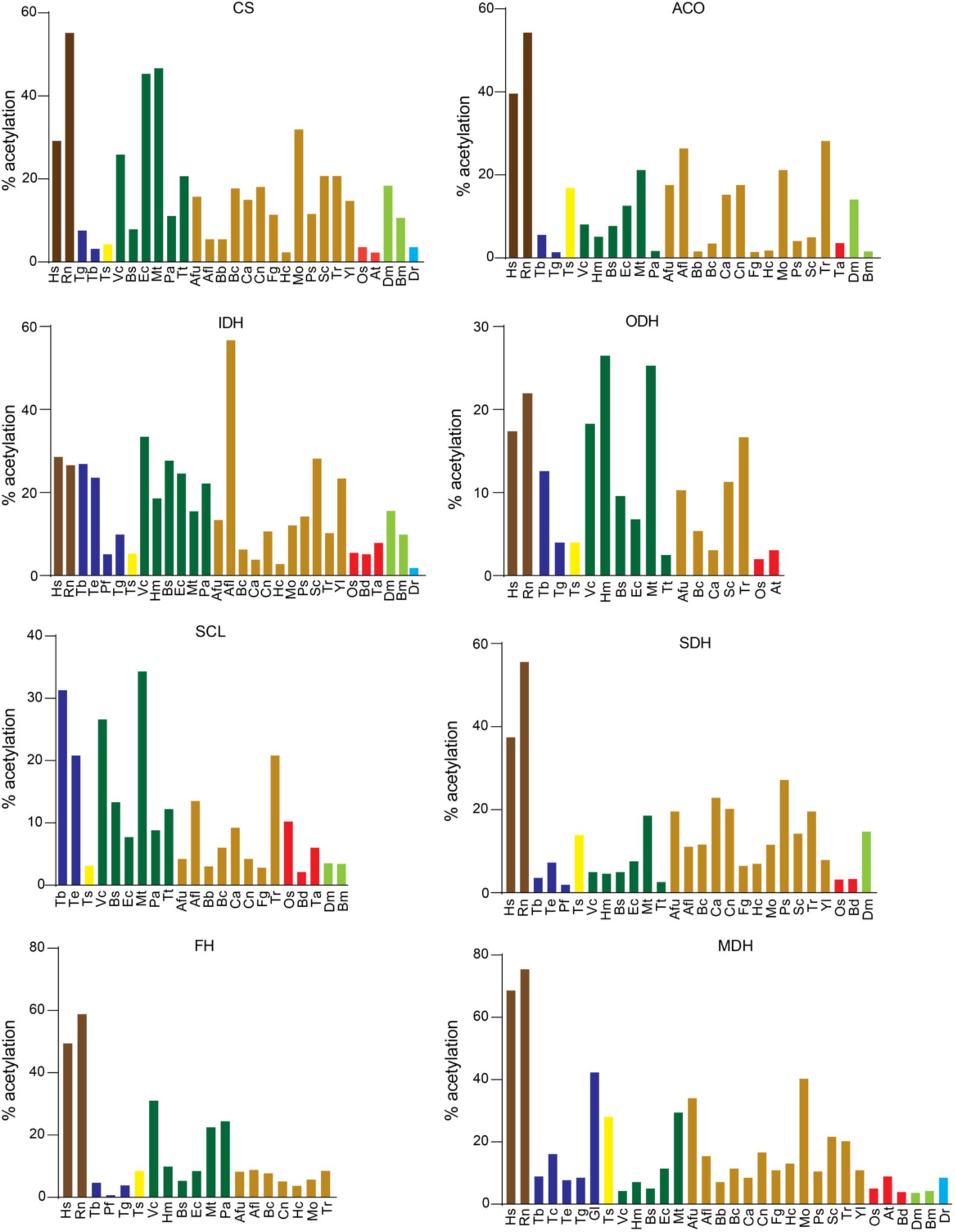
TCA cycle enzymes percentage of the lysine-acetylated and non-acetylated found in each group of organisms of different species. Enzymes were considered acetylated even if only one Kac site had been identified in the specific acetylome. Citrate synthase (CS), Aconitate (ACO), Isocitrate dehydrogenase (IDH), α-Ketoglutarate (ODH), Succinyl-CoA synthetase (SCL), Succinate dehydrogenase (SDH), Fumarate Hydratase (FH), Malate dehydrogenase (MDH). The groups are represented by different colors as follows: mammals (brown); protozoan (blue); worms (yellow); bacteria (green); fungi (gold); plants (red); insects (light green); fish (cyan). Identified species: Hs (*Homo sapiens)*; Rn (*Rattus* novergicus); Tb (*trypanosoma* brucei); Tc (*Trypanosoma cruzi*); Tg (*Toxoplasma gondii*); Pf (*Plasmodium falciparum*); Te (*Trypanosoma evansi*); Gl (*Giardia lamblia*); Hm (*Haloferax mediterranei*); Bs (*Bacillus subtilis*); Ec (*Escherichia coli*); Mt (*Mycobacterium tuberculosis*); Pa (*Pseudomonas aeruginosa*); Tt (*Thermus thermophilus*); Vc (*Vibrio colerae*); Afl (*Aspergillus flavus*); Afu (*Aspergillus fumigatus*); Bb (*Beauveria bassiana*); Bc (*Botrytis cinerea*); Ca (*Candida albicans*); Cn (*Cryptococcus neoformans*); Fg (*Fusarium graminearium*); Hc (*Histoplasma capsulatum*); Mo (*Magnaporthe oryzae*); Ps (*Phytophthora sojae*); Sc (*Saccharomyces cerevisiae*); Tr (*Trichophyton rubrum*); Yl (*Yarrowia lipolytica*); At (*Arabdopsis thaliana*); Os (*Oryza sativa*); Ta (*Triticum aestivum*); Bd (*Brachypodium distachyon*); Vv (*Vitis vinifera*); Sj (*Schistosoma japonicum*); Ts (*Trichinella spiralis*); Bm (*Bombyx mori*); Dm (*Drosophila melanogaster*); Dr (*Danio rerio*).

**Figure S9.**
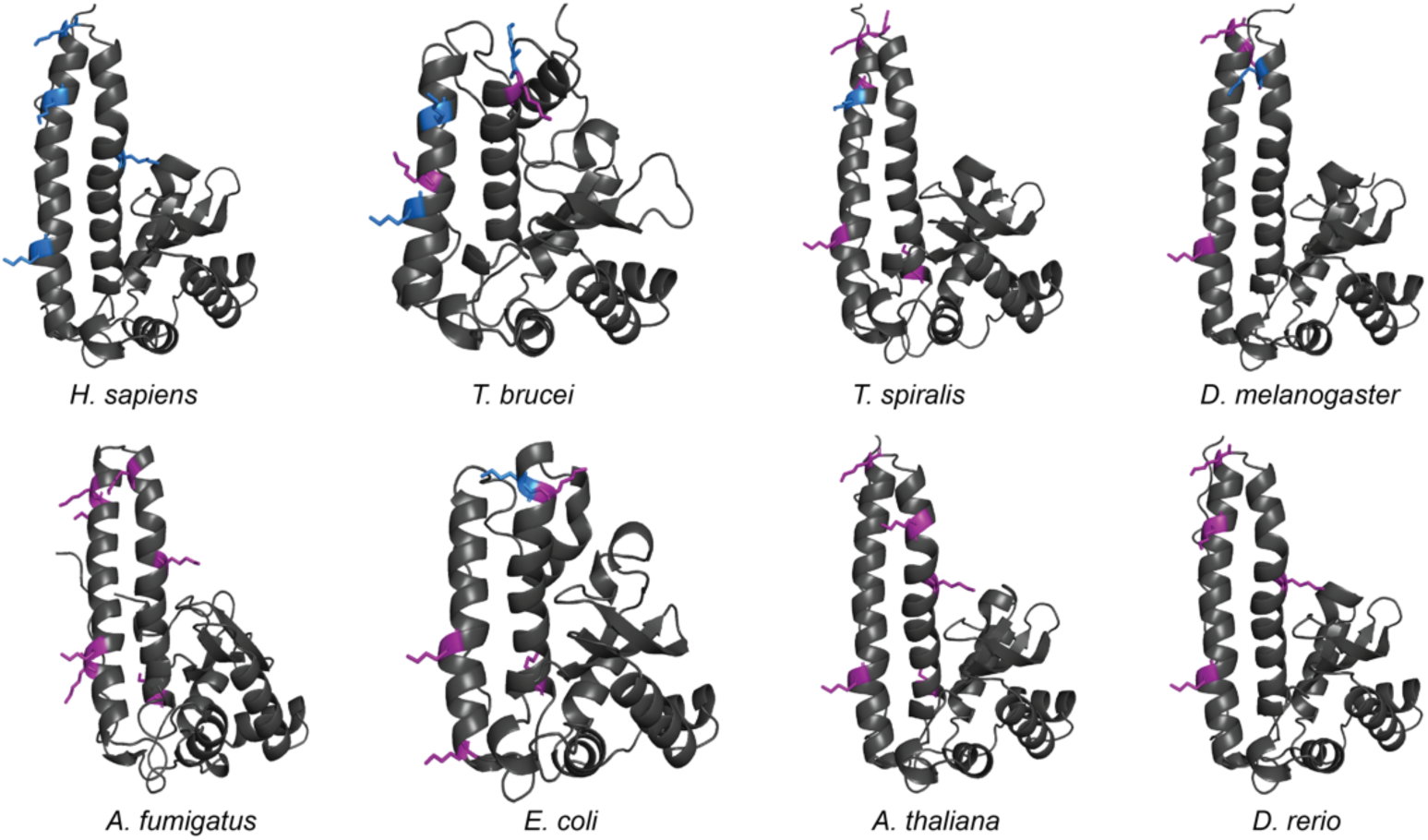
Lysine acetylation of the superoxide dismutase A across different groups. Highlighted within these structures are the lysine (K) residues identified as acetylated in their corresponding acetylomes (blue). Notably, several of these acetylated residues are located within the enzyme’s funnel region, which plays a crucial role in directing substrates towards the catalytic site. Blue: lysine residues found acetylated. Purple: lysine residues not yet detected acetylated.

**Supplementary Table 1.** List of selected acetylomes in this study.

| Group | Specie | Number of Kac sites | Number of Kac proteins | REF |
| --- | --- | --- | --- | --- |
| Archaea | *Haloferax mediterranei* | 1017 | 643 | Liu et al., 2018 |
| Bacteria | *Bacillus subtilis* | 1355 | 629 | Kosono et al., 2015 |
| Bacteria | *Escherichia coli* | 2502 | 809 | Castaño-Cerezo et al., 2014 |
| Bacteria | *Mycobacterium tuberculosis* | 1128 | 658 | Xie et al., 2015 |
| Bacteria | *Pseudomonas aeruginosa* | 1102 | 522 | Gaviard et al 2018 |
| Bacteria | *Thermus thermophilus* | 197 | 124 | Okanish et al., 2013 |
| Bacteria | *Vibrio cholerae* | 3402 | 1240 | Jers et al., 2018 |
| Fungi | *Aspergillus flavus* | 1383 | 652 | Lv et al., 2017 |
| Fungi | *Aspergillus fumigatus* | 5328 | 2312 | Li et al., 2019 |
| Fungi | *Beauveria bassiana* | 3421 | 1458 | Cai et al., 2021 |
| Fungi | *Botrytis cinerea* | 1582 | 954 | Lv et al., 2016 |
| Fungi | *Candida albicans* | 2048 | 926 | Li et al., 2019 |
| Fungi | *Cryptococcus neoformans* | 3535 | 1461 | Li et al., 2019 |
| Fungi | *Fusarium graminearum* | 517 | 364 | Zhou et al., 2016 |
| Fungi | *Histoplasma capsulatum* | 775 | 456 | Xie et al., 2016 |
| Fungi | *Magnaporthe oryzae hyphae* | 2720 | 1269 | Sun et al., 2017 |
| Fungi | *Magnaporthe oryzae mycelia* | 1551 | 704 | Liang et al 2018 |
| Fungi | *Phytophthora sojae* | 2197 | 1150 | Li et al., 2016 |
| Fungi | *Saccharomyces cerevisiae* | 4120 | 1321 | Weinert et al., 2014 |
| Fungi | *Trichophyton rubrum conidia* | 386 | 285 | Xu et al., 2018 |
| Fungi | *Trichophyton rubrum mycelia* | 5414 | 2335 | Xu et al., 2018 |
| Fungi | *Yarrowia lipolytica* | 3163 | 1428 | Wang et al., 2017 |
| Protozoan | *Plasmodium falciparum* | 2876 | 1146 | Cobbold et al., 2016 |
| Protozoan | *Toxoplasma gondii* (extra) | 571 | 386 | Xue et al., 2013 |
| Protozoan | *Toxoplasma gondii* (intra) | 411 | 274 | Jeffers et al., 2012 |
| Protozoan | *Trypanosoma brucei* (blood) |  |  | Zhang et al., 2020 |
| Protozoan | *Trypanosoma brucei* (procyclic) | 288 | 210 | Moretti et al 2018 |
| Protozoan | *Trypanosoma evansi* |  |  | Zhang et al., 2020 |
| Protozoan | *Trypanosoma cruzi* | 389 | 235 | Moretti et al 2018 |
| Protozoan | *Giardia lamblia* | 2999 | 956 | Zhu et al., 2021 |
| Worms | *Schistosoma japonicum* (adult) | 2393 | 1109 | Hong et al 2016 |
| Worms | *Schistosoma japonicum* (juvenile male) | 102 | 71 | Li et al., 2017 |
| Worms | *Schistosoma japonicum* (juvenile female) | 231 | 162 | Li et al., 2017 |
| Worms | *Schistosoma japonicum* (adult male) | 616 | 346 | Li et al., 2017 |
| Worms | *Schistosoma japonicum* (adult female) | 430 | 281 | Li et al., 2017 |
| Worms | *Trichinella spiralis* | 3872 | 1592 | Yang et al., 2018 |
| Plants | *Arabidopsis thaliana* | 2152 | 1022 | Hartl et al., 2017 |
| Plants | *Brachypodium distachyon* | 636 | 353 | Zhen et al 2016 |
| Plants | *Fragaria* sp | 1392 | 684 | Fang et al 2015 |
| Plants | *Oryza sativa* | 1353 | 866 | Xue et al., 2020 |
| Plants | *Triticum aestivum* | 416 | 277 | Zhang et al., 2016 |
| Insects | *Bombyx mori* | 667 | 342 | Nie et al., 2015 |
| Insects | *Drosophila melanogaster* | 1981 | 1013 | Weinert et al 2011 |
| Fish | *Danio rerio* | 377 | 189 | Kwon et al., 2016 |
| Mammals | HeLA cells | 3345 | 1440 | Xu et al., 2016 |
| Mammals | U2OS cells | 3174 | 855 | Sol et al., 2012 |
| Mammals | Human skeletal muscle tissue | 2811 | 991 | Lundby et al., 2012 |
| Mammals | *Rattus norvegicus* (16 different tissues) | 15474 | 4541 | Lundby et al., 2012 |

## REFERENCES

1 Ramazi, S., Allahverdi, A. and Zahiri, J. (2020) Evaluation of post-translational modifications in histone proteins: A review on histone modification defects in developmental and neurological disorders. J Biosci 10.1007/s12038-020-00099-2

2 Ryšlavá, H., Doubnerová, V., Kavan, D. and Vaněk, O. (2013) Effect of posttranslational modifications on enzyme function and assembly. J Proteomics 10.1016/j.jprot.2013.03.025

3 Mann, M. and Jensen, O. N. (2003) Proteomic analysis of post-translational modifications. Nat Biotechnol 10.1038/nbt0303-255

4 Phillips, D. M. (1963) The presence of acetyl groups of histones. Biochem J 87 10.1042/bj0870258

5 Ali, I., Conrad, R. J., Verdin, E. and Ott, M. (2018) Lysine Acetylation Goes Global: From Epigenetics to Metabolism and Therapeutics. Chem Rev 10.1021/acs.chemrev.7b00181

6 Choudhary, C., Weinert, B. T., Nishida, Y., Verdin, E. and Mann, M. (2014) The growing landscape of lysine acetylation links metabolism and cell signalling. Nat Rev Mol Cell Biol 10.1038/nrm3841

7 Narita, T., Weinert, B. T. and Choudhary, C. (2019) Functions and mechanisms of non-histone protein acetylation. Nat Rev Mol Cell Biol 10.1038/s41580-018-0081-3

8 Choudhary, C., Kumar, C., Gnad, F., Nielsen, M. L., Rehman, M., Walther, T. C., et al. (2009) Lysine acetylation targets protein complexes and co-regulates major cellular functions. Science (1979) 325 10.1126/science.1175371

9 Weinert, B. T., Iesmantavicius, V., Moustafa, T., Schölz, C., Wagner, S. A., Magnes, C., et al. (2014) Acetylation dynamics and stoichiometry in Saccharomyces cerevisiae. Mol Syst Biol 10 10.1002/msb.134766

10 Weinert, B. T., Wagner, S. A., Horn, H., Henriksen, P., Liu, W. R., Olsen, J. V., et al. (2011) Proteome-wide mapping of the Drosophila acetylome demonstrates a high degree of conservation of lysine acetylation. Sci Signal 4 10.1126/scisignal.2001902

11 Castaño-Cerezo, S., Bernal, V., Post, H., Fuhrer, T., Cappadona, S., Sánchez-Díaz, N. C., et al. (2014) Protein acetylation affects acetate metabolism, motility and acid stress response in Escherichia coli . Mol Syst Biol 10 10.15252/msb.20145227

12 Xue, B., Jeffers, V., Sullivan, W. J. and Uversky, V. N. (2013) Protein intrinsic disorder in the acetylome of intracellular and extracellular Toxoplasma gondii. Mol Biosyst 9 10.1039/c3mb25517d

13 Jeffers, V. and Sullivan, W. J. (2012) Lysine acetylation is widespread on proteins of diverse function and localization in the protozoan parasite Toxoplasma gondii. Eukaryot Cell 11 10.1128/EC.00088-12

14 Cobbold, S. A., Santos, J. M., Ochoa, A., Perlman, D. H. and Llinas, M. (2016) Proteome-wide analysis reveals widespread lysine acetylation of major protein complexes in the malaria parasite. Sci Rep 6 10.1038/srep19722

15 Nakayasu, E. S., Burnet, M. C., Walukiewicz, H. E., Wilkins, C. S., Shukla, A. K., Brooks, S., et al. (2017) Ancient regulatory role of lysine acetylation in central metabolism. mBio 8 10.1128/mBio.01894-17

16 Shan, Q., Ma, F., Wei, J., Li, H., Ma, H. and Sun, P. (2019) Physiological Functions of Heat Shock Proteins. Curr Protein Pept Sci 21 10.2174/1389203720666191111113726

17 Seo, J. H., Park, J. H., Lee, E. J., Vo, T. T. L., Choi, H., Kim, J. Y., et al. (2016) ARD1-mediated Hsp70 acetylation balances stress-induced protein refolding and degradation. Nat Commun 7 10.1038/ncomms12882

18 Lundby, A., Lage, K., Weinert, B. T., Bekker-Jensen, D. B., Secher, A., Skovgaard, T., et al. (2012) Proteomic Analysis of Lysine Acetylation Sites in Rat Tissues Reveals Organ Specificity and Subcellular Patterns. Cell Rep 2 10.1016/j.celrep.2012.07.006

19 Zhao, S., Xu, W., Jiang, W., Yu, W., Lin, Y., Zhang, T., et al. (2010) Regulation of cellular metabolism by protein lysine acetylation. Science (1979) **327** 10.1126/science.1179689

20 Yang, L., Vaitheesvaran, B., Hartil, K., Robinson, A. J., Hoopmann, M. R., Eng, J. K., et al. (2011) The fasted/fed mouse metabolic acetylome: N6-acetylation differences suggest acetylation coordinates organ-specific fuel switching. J Proteome Res 10 10.1021/pr200313x

21 Wang, Q., Zhang, Y., Yang, C., Xiong, H., Lin, Y., Yao, J., et al. (2010) Acetylation of metabolic enzymes coordinates carbon source utilization and metabolic flux. Science (1979*<otherinfo>)* 327**</otherinfo>** 10.1126/science.1179687

22 Gamblin, S. J., Davies, G. J., Grimes, J. M., Jackson, R. M., Littlechild, J. A. and Watson, H. C. (1991) Activity and specificity of human aldolases. J Mol Biol 219 10.1016/0022-2836(91)90650-U

23 Leite, A. B., Gomes, A. A. S., Sousa, A. C. D. C. N., Fontes, M. R. D. M., Schenkman, S. and Moretti, N. S. (2020) Effect of lysine acetylation on the regulation of Trypanosoma brucei glycosomal aldolase activity. Biochemical Journal 477 10.1042/BCJ20200142

24 Kang, W., Suzuki, M., Saito, T. and Miyado, K. (2021) Emerging role of tca cycle-related enzymes in human diseases. Int J Mol Sci 10.3390/ijms222313057

25 Broxton, C. N. and Culotta, V. C. (2016) SOD Enzymes and Microbial Pathogens: Surviving the Oxidative Storm of Infection. PLoS Pathog 10.1371/journal.ppat.1005295

26 dos Santos Moura, L., Santana Nunes, V., Gomes, A. A. S., Sousa, A. C. de C. N., Fontes, M. R. M., Schenkman, S., et al. (2021) Mitochondrial Sirtuin TcSir2rp3 Affects TcSODA Activity and Oxidative Stress Response in Trypanosoma cruzi. Front Cell Infect Microbiol 11 10.3389/fcimb.2021.773410

27 Lu, J., Cheng, K., Zhang, B., Xu, H., Cao, Y., Guo, F., et al. (2015) Novel mechanisms for superoxide-scavenging activity of human manganese superoxide dismutase determined by the K68 key acetylation site. Free Radic Biol Med 85 10.1016/j.freeradbiomed.2015.04.011

28 Maran, S. R., Fleck, K., Monteiro-Teles, N. M., Isebe, T., Walrad, P., Jeffers, V., et al. (2021) Protein acetylation in the critical biological processes in protozoan parasites. Trends Parasitol 37, 815–830 10.1016/j.pt.2021.04.008

29 Mayer, M. P. and Bukau, B. (2005) Hsp70 chaperones: Cellular functions and molecular mechanism. Cellular and Molecular Life Sciences 10.1007/s00018-004-4464-6

30 Rosenzweig, R., Nillegoda, N. B., Mayer, M. P. and Bukau, B. (2019) The Hsp70 chaperone network. Nat Rev Mol Cell Biol 10.1038/s41580-019-0133-3

31 Kaushik, S. and Cuervo, A. M. (2018) The coming of age of chaperone-mediated autophagy. Nat Rev Mol Cell Biol 10.1038/s41580-018-0001-6

32 Griffith, A. A. and Holmes, W. (2019) Fine tuning: Effects of post-translational modification on Hsp70 chaperones. Int J Mol Sci 10.3390/ijms20174207

33 Nitika and Truman, A. W. (2017) Cracking the Chaperone Code: Cellular Roles for Hsp70 Phosphorylation. Trends Biochem Sci 10.1016/j.tibs.2017.10.002

34 Cloutier, P. and Coulombe, B. (2013) Regulation of molecular chaperones through post-translational modifications: Decrypting the chaperone code. Biochim Biophys Acta Gene Regul Mech 10.1016/j.bbagrm.2013.02.010

35 Nitika, Porter, C. M., Truman, A. W. and Truttmann, M. C. (2020) Post-translational modifications of Hsp70 family proteins: Expanding the chaperone code. Journal of Biological Chemistry 10.1074/jbc.REV120.011666

36 Marmorstein, R. and Zhou, M.-M. (2014) Writers and readers of histone acetylation: structure, mechanism, and inhibition. Cold Spring Harb Perspect Biol 6, a018762 10.1101/cshperspect.a018762

37 Shvedunova, M. and Akhtar, A. (2022) Modulation of cellular processes by histone and non-histone protein acetylation. Nat Rev Mol Cell Biol 10.1038/s41580-021-00441-y

38 Penkler, G., Du Toit, F., Adams, W., Rautenbach, M., Palm, D. C., Van Niekerk, D. D., et al. (2015) Construction and validation of a detailed kinetic model of glycolysis in Plasmodium falciparum. FEBS Journal 282 10.1111/febs.13237

39 Bei, J., Chen, Y., Zhang, Q., Wang, X., Lin, L., Huang, J., et al. (2023) HBV suppresses macrophage immune responses by impairing the TCA cycle through the induction of CS/PDHC hyperacetylation. Hepatol Commun 10.1097/HC9.0000000000000294

40 Bradshaw, P. C. (2021) Acetyl-coa metabolism and histone acetylation in the regulation of aging and lifespan. Antioxidants 10 10.3390/antiox10040572

41 Hu, M., You, Y., Li, Y., Ma, S., Li, J., Miao, M., et al. (2023) Deacetylation of ACO2 Is Essential for Inhibiting Bombyx mori Nucleopolyhedrovirus Propagation. Viruses 15 10.3390/v15102084

42 Balparda, M., Elsässer, M., Badia, M. B., Giese, J., Bovdilova, A., Hüdig, M., et al. (2022) Acetylation of conserved lysines fine-tunes mitochondrial malate dehydrogenase activity in land plants. Plant Journal 109 10.1111/tpj.15556

43 Venkat, S., Gregory, C., Sturges, J., Gan, Q. and Fan, C. (2017) Studying the Lysine Acetylation of Malate Dehydrogenase. J Mol Biol 429 10.1016/j.jmb.2017.03.027

44 Fridovich, I. (1983) Superoxide radical: An endogenous toxicant. Annu Rev Pharmacol Toxicol **Vol.** 23 10.1146/annurev.pa.23.040183.001323

45 Benovic, J., Tillman, T., Cudd, A. and Fridovich, I. (1983) Electrostatic facilitation of the reaction catalyzed by the manganese-containing and the iron-containing superoxide dismutases. Arch Biochem Biophys 221 10.1016/0003-9861(83)90151-0

46 Soppa, J. (2010) Protein acetylation in archaea, bacteria, and eukaryotes. Archaea 10.1155/2010/820681

47 Jumper, J., Evans, R., Pritzel, A., Green, T., Figurnov, M., Ronneberger, O., et al. (2021) Highly accurate protein structure prediction with AlphaFold. Nature 596 10.1038/s41586-021-03819-2

48 Waterhouse, A., Bertoni, M., Bienert, S., Studer, G., Tauriello, G., Gumienny, R., et al. (2018) SWISS-MODEL: Homology modelling of protein structures and complexes. Nucleic Acids Res 46 10.1093/nar/gky427

49 Pronk, S., Páll, S., Schulz, R., Larsson, P., Bjelkmar, P., Apostolov, R., et al. (2013) GROMACS 4.5: A high-throughput and highly parallel open source molecular simulation toolkit. Bioinformatics 29 10.1093/bioinformatics/btt055

50 Huang, J., Rauscher, S., Nawrocki, G., Ran, T., Feig, M., De Groot, B. L., et al. (2016) CHARMM36m: An improved force field for folded and intrinsically disordered proteins. Nat Methods 14 10.1038/nmeth.4067

51 Warnecke, A., Sandalova, T., Achour, A. and Harris, R. A. (2014) PyTMs: A useful PyMOL plugin for modeling common post-translational modifications. BMC Bioinformatics 15 10.1186/s12859-014-0370-6

52 Bussi, G., Donadio, D. and Parrinello, M. (2007) Canonical sampling through velocity rescaling. Journal of Chemical Physics 126 10.1063/1.2408420

53 Eslami, H., Mojahedi, F. and Moghadasi, J. (2010) Molecular dynamics simulation with weak coupling to heat and material baths. Journal of Chemical Physics 133 10.1063/1.3474951

54 Hoover, W. G. (1985) Canonical dynamics: Equilibrium phase-space distributions. Phys Rev A (Coll Park) 31 10.1103/PhysRevA.31.1695

55 Nosé, S. and Klein, M. L. (1986) Constant-temperature-constant-pressure molecular-dynamics calculations for molecular solids: Application to solid nitrogen at high pressure. Phys Rev B 33 10.1103/PhysRevB.33.339

56 Laurent, B., Chavent, M., Cragnolini, T., Dahl, A. C. E., Pasquali, S., Derreumaux, P., et al. (2015) Epock: Rapid analysis of protein pocket dynamics. Bioinformatics 31 10.1093/bioinformatics/btu822

57 Humphrey, W., Dalke, A. and Schulten, K. (1996) VMD: Visual molecular dynamics. J Mol Graph 14 10.1016/0263-7855(96)00018-5

